# Heat tolerance of tropical herbaceous plants increases with elevation

**DOI:** 10.1101/2025.03.12.642681

**Authors:** Georgia G. Hernández, Jeffrey R. Seemann, Lucas da Cunha Godoy, Martijn Slot, Carlos García-Robledo

## Abstract

**Background and Aims:** Tropical plants are assumed to be especially vulnerable to global warming because their physiologies are adapted to relatively constant temperatures throughout the year. Furthermore, it has been found that woody plants in colder high elevation environments are less tolerant to high temperatures than plants in the warmer lowlands. Here, we examined heat tolerance in a group of herbaceous plants with a wide elevational distribution in the tropics.

**Methods:** This study focused on 61 species from the order Zingiberales (ginger and banana-like plants) distributed from the lowlands (50 m asl) to lower montane forests (2000 m asl) along the Barva elevational gradient in Costa Rica.

This study addressed the following questions: **a)** Does heat tolerance of Zingiberales species differ along the elevational gradient? **b)** Does heat tolerance vary along the elevational gradient within families of Zingiberales? **c)** Does heat tolerance vary along the elevational gradient within species for those with broad elevational distributions? To test if the temperature that causes damage to the function of photosystem II (PSII) in Zingiberales is associated with the temperatures prevalent at their elevation, we estimated heat tolerance (T_50_) of PSII using chlorophyll fluorescence techniques.

**Key Results:** In contrast to the results found in tropical trees, our results showed that T_50_ is higher at higher elevations than in the lowlands for herbaceous plants species. This trend was observed across plant communities and families, and within most species with wide distributions along the elevational gradient.

**Conclusions:** Our study suggests that herbs differ from trees in their elevational patterns in heat tolerance. We hypothesize that maximum and minimum leaf temperatures, and UV radiation may play a role in the observed pattern.

## INTRODUCTION

The continuing increase in global temperature predicted for the current century (between 0.9 - 5.3°C; Lee et al. 2021) is also expected to increase the risks of thermal stress in terrestrial vegetation, including tropical plant species (Doughty et al. 2023; Mau et al. 2018). Field observations have already shown that tropical plants are particularly sensitive to small increases in temperature that can reduce tree growth (Clark et al., 2003; Feeley et al., 2007) and photosynthesis (Doughty and Goulden 2008), and increase mortality (Clark et al., 2010). For tropical tree species the temperature range over which they can maintain maximum rates of photosynthesis is narrower than temperate tree species, which can make them more vulnerable to predicted increases in global temperatures (Cunningham and Read 2002). Plant species are generally adapted to their thermal environments (Kumarathunge et al. 2019). It has been assumed that given the relatively low variation in temperature at any particular elevation within tropical mountains, organisms will be selected to have narrow thermal limits at each particular elevation (Addo-Bediako, Chown, and Gaston 2000; Janzen 1967; Polato et al. 2018; Sunday, Bates, and Dulvy 2012). As temperatures changes due to global warming, plants must locally adapt or migrate to avoid heat damage (Feeley et al. 2023; Holt 1990; Malhi et al. 2010). Understanding how photosynthetic thermal limits change along a tropical mountain elevational gradient is important in determining the thermal plasticity and the need for adaptation or migration for plants at different elevations.

Plant species with broad distributions often exhibit great adaptability to survive and thrive across diverse environments (Joshi et al. 2001). To cope with local conditions, these species may rely on ecotypic differentiation (Leimu and Fischer 2008; Wadgymar, Daws, and Anderson 2017). Ecotypes represent locally adapted populations with distinct physiological characteristics that may enhance their fitness within their native environments (McGraw and Antonovics 1983). Local adaptation, such as ecotypic differentiation along elevational gradients (McGraw and Antonovics 1983)., can confer a competitive advantage, enabling species to expand their geographic range. Although plasticity benefits some plant species, genetic differentiation is essential for others to persist across steep environmental gradients. In a rapidly changing climate, both phenotypic plasticity and genetic adaptation will be key determinants of a plant species’ ability to persist (Nicotra et al. 2015; Pfennigwerth, Bailey, and Schweitzer 2017).

Tropical elevational gradients are natural laboratories for studying organismal responses to global warming (Malhi et al. 2010). In tropical mountains, temperature decreases by 0.5 - 0.6°C with an increase of 100 m in elevation (D. Clark, Hurtado, and Saatchi 2015; McCain and Grytnes 2010). If thermal niches of tropical plants are narrow, a small increase in temperature will exert a significant thermal stress on plants’ photosynthetic machinery, and ultimately growth and survival. One parameter used to measure plant stress is the temperature at which the potential efficiency of photosynthetic system II (PSII) is reduced by 50% (T_50_) (Knight and Ackerly 2002; Krause et al. 2010).

Our study site is the Barva elevational transect in Costa Rica, a continuous forested area that extends from lowland tropical wet forest (50 m asl) to upper montane cloud forest (2830 m asl). The mean annual temperature for the Barva transect ranges from 25°C in the lowlands to 10.4°C at the highest elevation (D. Clark, Hurtado, and Saatchi 2015). Current evidence suggests that the mean annual temperature has been increasing at a rate of 0.017°C per year since 1950 at the lowland forest of La Selva Biological Station (hereafter, La Selva) (Feeley et al., 2013, Harris et al., 2020). This temperature change since 1950 is equivalent to an elevation shift of 200-240m. Long-term temperature records show that global warming already affects tropical biomes along this mountain, with species composition shifting along the Barva transect (Feeley et al., 2013). This upward migration and shift in composition has been suggested to be driven by mortality events on the warm edge of the distribution (Feeley et al., 2013), highlighting the importance of heat tolerance in evaluating climate change effects on plant populations.

In this study, we focus on herbaceous plants from the order Zingiberales, a charismatic group of tropical plants that includes the families Cannaceae, Costaceae, Heliconiaceae, Marantaceae, Musaceae, and Zingiberaceae. To determine their thermal limits, we estimated heat tolerance (T_50_) for 61 species distributed along the Barva transect. We addressed the following questions: **a)** Does heat tolerance of Zingiberales species differ along the elevational gradient? **b)** Does heat tolerance vary along the elevational gradient within families of Zingiberales? **c)** Does heat tolerances vary along the elevational gradient within species for those with broad elevational distributions? Although this study cannot definitively distinguish between acclimation and adaptation based solely on in situ observations (Halbritter et al. 2018), it aims to investigate the phenotypic variation within populations of a multiple species distributed across a broad tropical elevational gradient. We expect plants to be adapted to the prevailing temperature at each elevation. We also anticipate that populations will be locally adapted along the elevational gradient for those species with broad distributions, exhibiting higher heat tolerance in areas with the highest local temperatures.

## MATERIAL AND METHODS

### Study site

This study was conducted along the La Selva – Volcán Barva elevational gradient (hereafter the Barva transect) from 2017 to 2019, with measurements restricted to the wet season (May-August). We incorporated heat tolerance measurements for Zingiberales at 50 m asl from Hernández *et al*. (2022), collected at the La Selva. The Barva transect is located on the Caribbean slope of northeastern Costa Rica (10.28° - 10.18°N, 84.02° - 84.11°W). The Barva transect extends from the lowland tropical wet forest of La Selva Biological Station (50 m asl) to montane forests in the protected area of the Braulio Carrillo National Park (2830 m asl). The Barva is the largest elevational gradient of continuous forest in Central America, with an area of 50,000 ha within the boundaries of Braulio Carillo National Park and the La Selva Biological Station (D. Clark, Hurtado, and Saatchi 2015). Part of this study was also conducted at the Albergue El Socorro, a private forest adjacent to the Lower Premontane Rainforest in Braulio Carrillo National Park.

Annual rainfall varies along the Barva transect from 4000 mm in the lowlands, 9000 mm at intermediate elevations (D. Clark, Hurtado, and Saatchi 2015), and 3300 mm at the summit (Hartshorn and Peralta 1988). Mean annual temperatures along the Barva transect range from 25°C in the lowlands to 10.4°C at the highest elevation, decreasing by 0.5°C for every 100 m in elevation (D. Clark, Hurtado, and Saatchi 2015). Our study sites spanned three life zones and two transition zones: Wet Forest at 50 m asl, Wet Forest, cool transition at 500 m asl, Lower Premontane Rainforest at 1000 m asl, Higher Premontane Rainforest at 1500 m asl, and Lower Montane Rainforest at 2000 m asl (Table 1; D. Clark, Hurtado, and Saatchi 2015; Hartshorn and Peralta 1988).

**Table 1.**
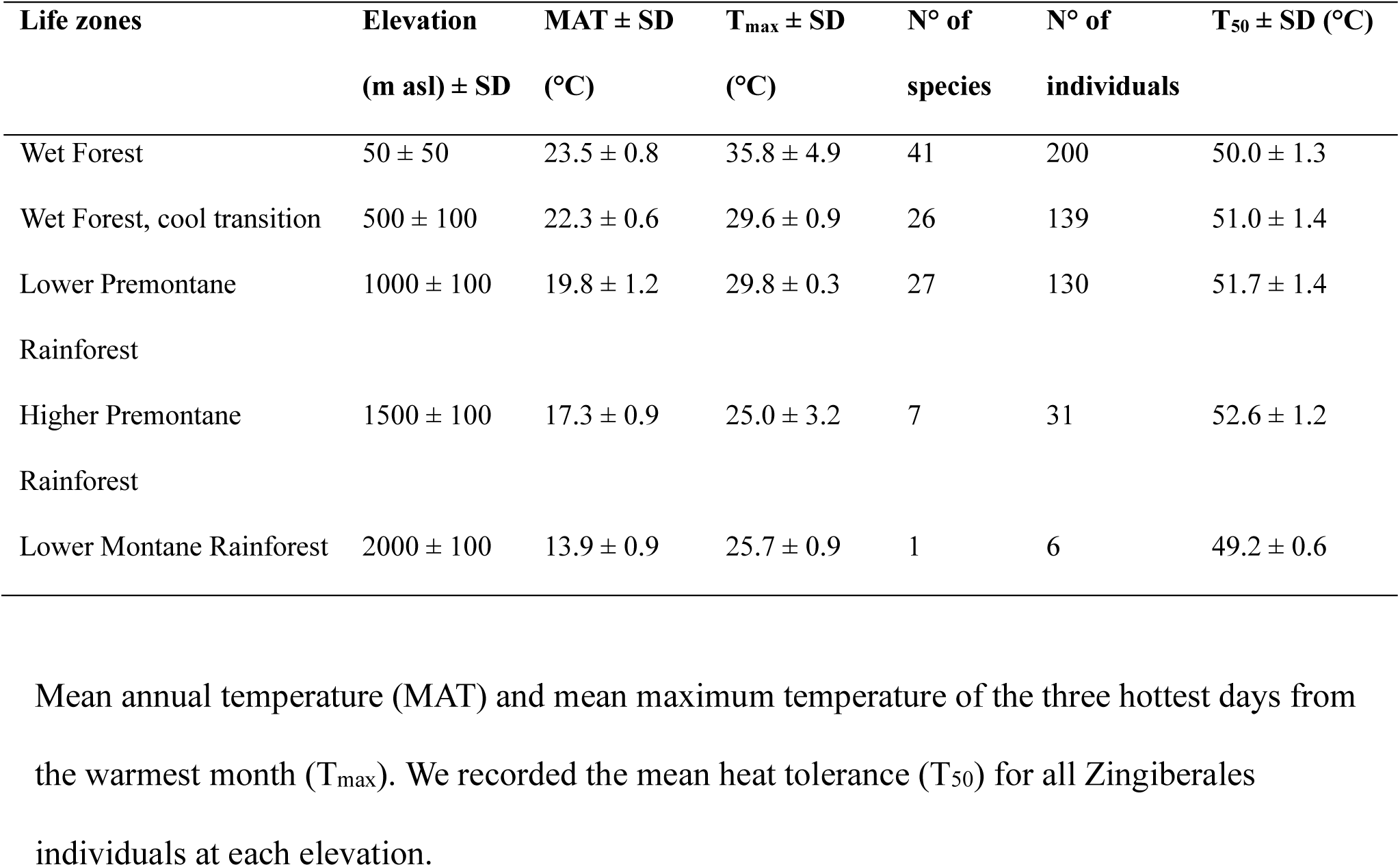
Life zone classifications as proposed by Hartshorn & Peralta (1988) and the elevational range of distribution where plants were collected.

| Life zones | Elevation<br>(m asl) ± SD | MAT ± SD<br>(°C) | T<sub>max</sub> ± SD<br>(°C) | N° of<br>species | N° of<br>individuals | T<sub>50</sub> ± SD (°C) |
| --- | --- | --- | --- | --- | --- | --- |
| Wet Forest | 50 ± 50 | 23.5 ± 0.8 | 35.8 ± 4.9 | 41 | 200 | 50.0 ± 1.3 |
| Wet Forest, cool transition | 500 ± 100 | 22.3 ± 0.6 | 29.6 ± 0.9 | 26 | 139 | 51.0 ± 1.4 |
| Lower Premontane<br>Rainforest | 1000 ± 100 | 19.8 ± 1.2 | 29.8 ± 0.3 | 27 | 130 | 51.7 ± 1.4 |
| Higher Premontane<br>Rainforest | 1500 ± 100 | 17.3 ± 0.9 | 25.0 ± 3.2 | 7 | 31 | 52.6 ± 1.2 |
| Lower Montane Rainforest | 2000 ± 100 | 13.9 ± 0.9 | 25.7 ± 0.9 | 1 | 6 | 49.2 ± 0.6 |

Mean annual temperature (MAT) and mean maximum temperature of the three hottest days from the warmest month (T<sub>max</sub>). We recorded the mean heat tolerance (T<sub>50</sub>) for all Zingiberales individuals at each elevation.

### Study species

The highly diverse plant order of Zingiberales comprises eight families, including bananas, gingers, heliconias, and relatives (Carlsen et al. 2018), all herbaceous in growth form. We estimated heat tolerance of 61 Zingiberales species from the families Cannaceae, Costaceae, Heliconiaceae, Marantaceae, Musaceae, and Zingiberaceae along the Barva elevational gradient (Supplementary Data Table S1). We selected this order because of its high diversity along the Barva transect, its high abundance, its importance in supporting ecological interactions, and easy identification. The number of Zingiberales species varied naturally across elevation sites, decreasing with increasing elevation (Table 1).

### Leaf heat tolerance (T_50_) of Zingiberales species along the Barva transect

To determine leaf heat tolerance, we collected one to five undamaged, fully expanded mature leaves per individual. Zingiberales species are predominantly understory herbs, but we also include species present in open areas and gaps in the forest (Supplementary Data Table S1, Fig. S3). To reduce intraspecific variation in chlorophyll fluorescence estimates due to light adaptation within species, we only sampled leaves from the most common habitat of each species, i.e., the leaves were always collected either in full sun or shade within one species (Supplementary Data Table S1, Fig. S3). This comprised 66% of the collected species exclusively found in shaded areas and 34% in sun areas. We estimated heat tolerance (T_50_) as the temperature that caused a 50% reduction of the initial ratio of variable over maximum fluorescence, F_v_/F_m_ (maximum quantum yield of the Photosystem II, PSII), following standard protocols (Krause et al. 2010; 2013). T_50_ reflects irreversible damage to the photosynthetic apparatus, attributed to a detachment of the light-harvesting antenna complex, resulting in a loss of PSII function and an increase in chlorophyll fluorescence (Kouřil et al. 2004). An estimation that corresponds to the inactivation of PSII, such as T_50_, is ecologically significant because it is considered a key indicator of high temperature stress (Baker and Rosenqvist 2004) and the most temperature sensitive component of the photosynthetic apparatus to high temperature (Downton, Berry, and Seemann 1984; Havaux 1993; Seemann, Downton, and Berry 1986).

We determined heat tolerance using the method described in Krause et al. (2010) and other recent tropical studies (Feeley et al. 2020; Hernández et al. 2022; Slot et al. 2021; Tiwari et al. 2021). *See* Supplementary Data Fig. S1 for experiment setup photos. We cut disks (diameter = 1.9 cm) from each leaf, avoiding middle veins and leaf edges. Leaf disks were assigned to each temperature treatment at random. Leaf disks were placed in Miracloth® (EMD Millipore Corp, Massachusetts, USA) fabric to prevent anaerobiosis (Krause et al. 2010). Leaf disks were then placed in waterproof Whirl-Pak® bags (Nasco, Wisconsin, USA) and completely immersed in one of several circulating hot water baths (ANOVA Sous Vide Precision Cooker A2.2-120V-US 2014, San Francisco, California, USA) for 15 minutes. We set water baths at the following temperatures: 24°C (ambient temperature, representing the control), 38, 42, 44, 46, 48, 50, 52, 54, or 60°C (Perez et al. 2021). The water baths have a temperature accuracy of ± 0.1°C, and water temperature constantly was checked with a long-stem thermometer (accuracy: ± 0.2°C). After 15 minutes, we removed the leaf disks from the water baths and placed them in petri dishes lined with moist paper towels for 24 hours at ambient temperature. After this recovery period, we dark-adapted each leaf disk for 20 minutes and then measured F_0_ and F_m_, the basal and maximum fluorescence yield, respectively, with a chlorophyll fluorometer (Model OS-30P, OptiSciences, New Hampshire, USA). From these measurements, F_v_/F_m_ is calculated as (F_m_-F_o_) /F_m_. T_50_ is the temperature that causes a 50% reduction in F_v_/F_m_ compared to the control value. We estimated T_50_ by calculating the logistic non-linear least squares model of F_v_/F_m_ vs temperature treatment.

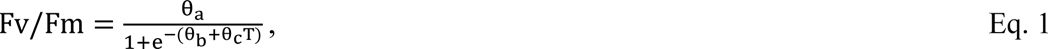

where T is the heat treatment temperature, θ_a_ is the asymptote of the heat treatment-response variable relationship, θ_b_ is a constant, and θ_c_ is the decay parameter. We fitted the logistic decay curve to pooled measurements per plant, with six leaf replicates at each temperature for each individual. Subsequently, we obtained the curve parameters per individual and the respective confidence intervals. We also obtanied the species-specific mean and standard deviation from the 5-replicate individual T_50_ values (Supplementary Data Table S1). The model was fitted with the “nls” function in the ‘stats’ package in R, version 4.0.4 (R Core Development Team 2024).

### Ambient temperature along the Barva transect

Temperature data were provided by D.B. Clark, D.A. Clark, TEAM project, and Conservation International (D. Clark, Hurtado, and Saatchi 2015). Data were obtained by placing one temperature sensor (Thermochron iButton® DS1921G-F5, resolution 0.5°C, Maxim Technology, San Jose, California) 1.5 m above ground in the understory of each of the elevation sites, from 2007 to 2011.

### Statistical analyses

We analyzed the relationship between heat tolerance and elevation using a linear model. To assess how heat tolerance is related to elevation and plant family, we fitted a linear mixed-effect model using the “lme4” package in R (R Core Development Team 2024). In the model, family, elevation, and their interaction were set as fixed effects with an interaction, while species was set as nested random within family. Most Zingiberales families have only a single genus, while Zingiberaceae and Marantaceae typically have one or two major genera (Table S1). Given this uneven representation of genera within families, we limited our analysis to the family and species levels. We estimated 95% confidence intervals of the linear mixed model after generating bootstrap means with 1000 iterations, using the “bootMer” function in the lme4 package (Bates et al. 2014).

We used a linear model to evaluate the relationship between heat tolerance and elevation within species. In this model, the interaction of elevation and species was set as a fixed effect. All models were subjected to residual checks for normality and homoscedasticity. All analyses were performed in R, version 4.0.4 (R Core Development Team 2024).

## RESULTS

We determined heat tolerance along the elevational gradient for 506 individuals representing 61 species, 17 genera, and 6 families in the order Zingiberales (Table 1). To test how species are adapted to temperature at different elevations along the Barva transect, we estimated the heat tolerance of Zingiberales species (Fig. 1). In disagreement with our initial predictions, the photosynthetic heat tolerance (T_50_) consistently increased with increasing elevation until the values steeply dropped when reaching the lower montane rainforest species at 2000 m asl (Fig. 1).

**Figure 1.**
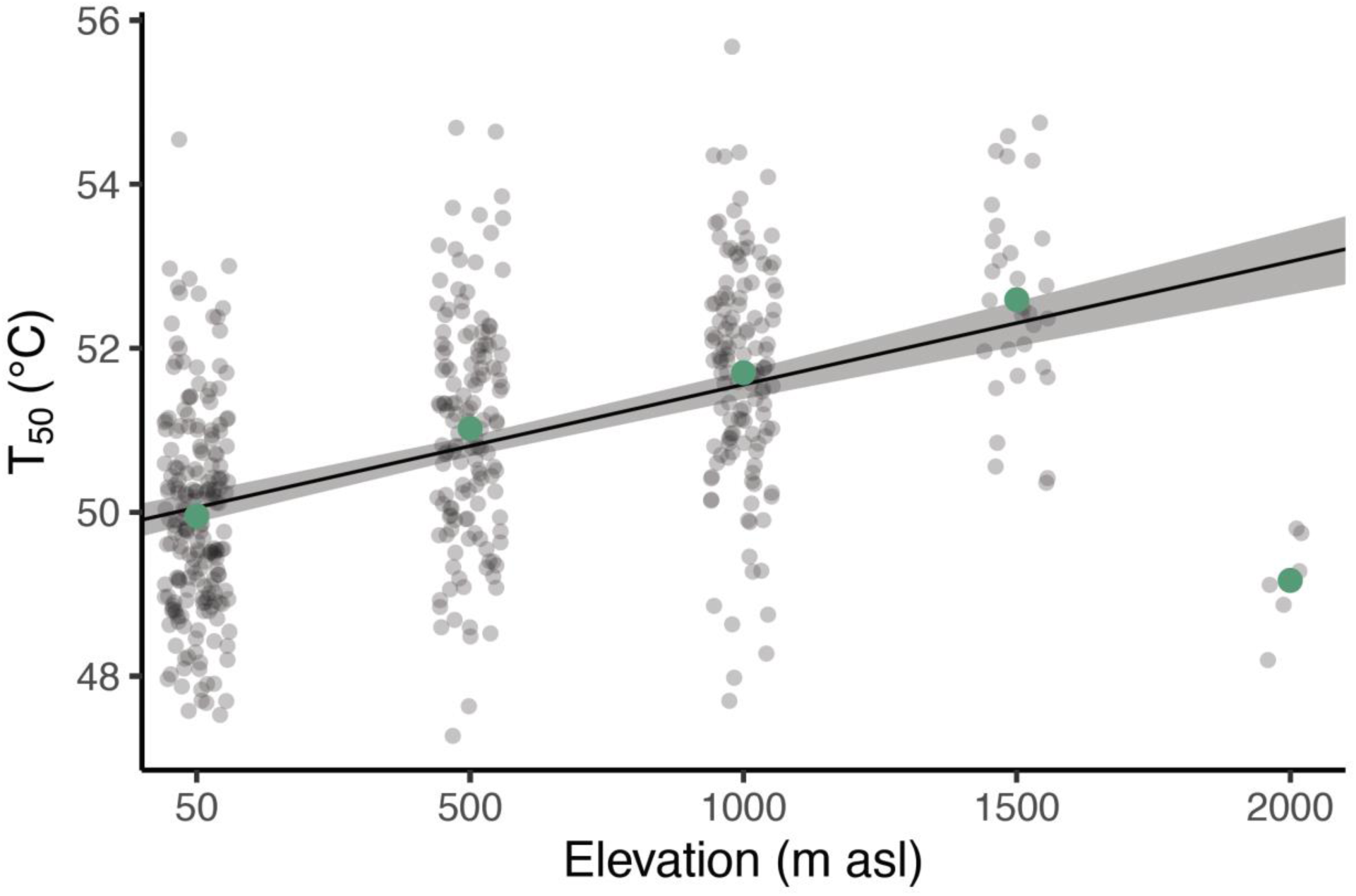
Relationship between heat tolerance of Zingiberales individuals and elevation at the Barva transect. Heat tolerance (T_50_) was estimated as the temperature at which PSII quantum yield (fluorescence) decreases by 50%. The line indicates linear regression from the model. Dots represent heat tolerance of individuals from each species at the given elevation site; the green dots represent the mean heat tolerance for each elevation. Shaded areas indicate 95% confidence intervals of the predictions. Note that at the highest elevation only one species is present. T_50_ = 50.06 + 0.0015 × Elevation, R^2^ = 0.21, P = < 0.0001.

Individuals sampled from the lowlands tended to have lower T_50_ than individuals from higher elevations (T_50_ = 50.1 + 0.0015 × Elevation, F = 135.31, R^2^ = 0.21, P < 2.2E-16). T_50_ increased by 0.15°C for every 100 m increase in elevation. However, elevation only explained a relatively low amount of variance of T_50_ (R^2^ = 0.21), as values varied considerably within each site (Fig. 1 and 2). The estimated means and standard deviation of T_50_ for all individuals at each elevation are shown in Table 1. Heat tolerance and the temperature range at each site of the Barva gradient are shown in Supplementary Data Fig. S5.

**Figure 2.**
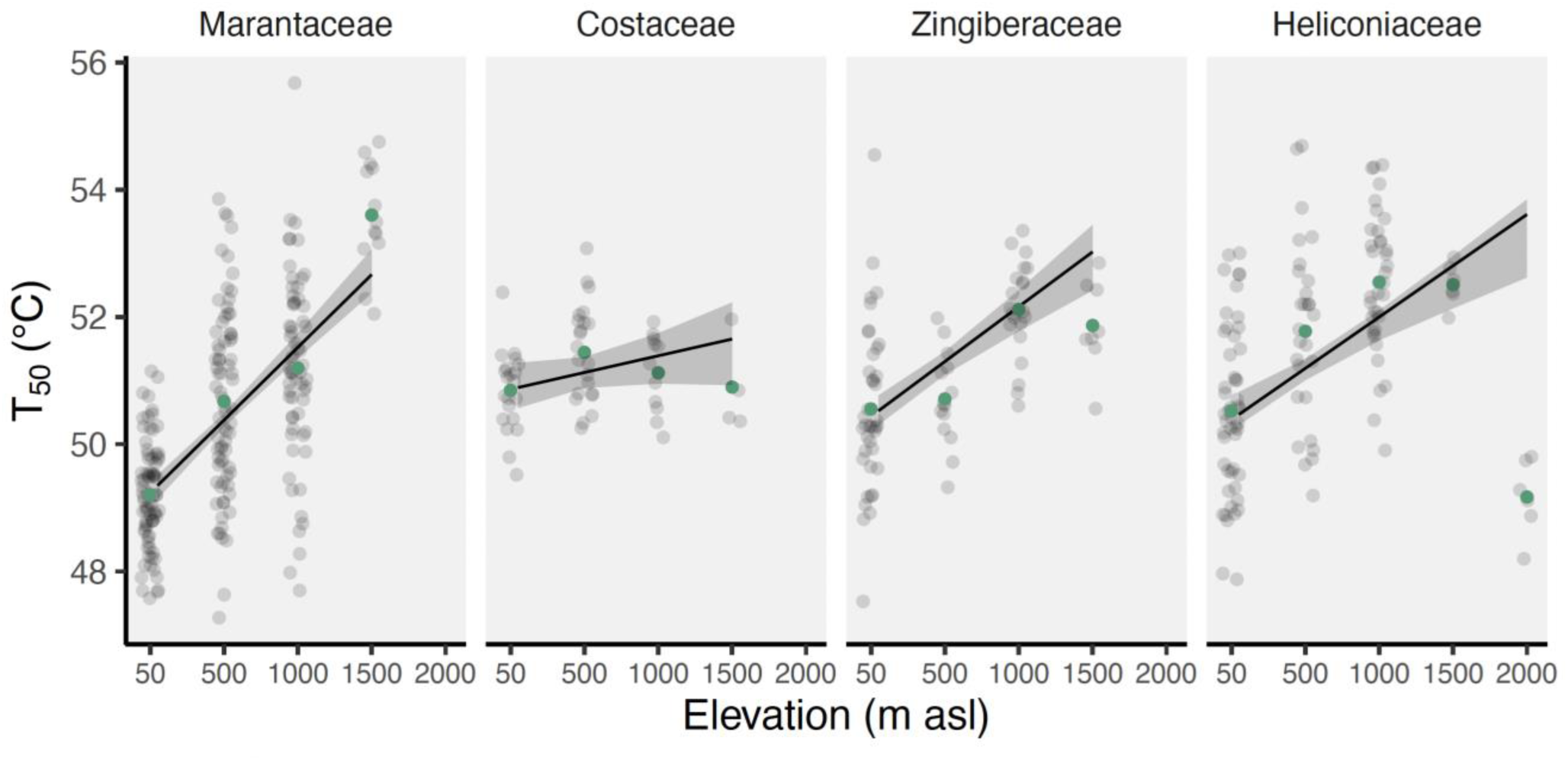
Relationship between heat tolerance (T_50_) of Zingiberales families and elevation at the Barva transect. Plant families are placed in their phylogenetic order. Lines indicate linear regression from the model. Dots represent heat tolerance of individual plants; the green dots represent the mean heat tolerance for each family at the given elevation. Shaded areas indicate 95% confidence intervals of the predictions. *See* Supplementary Data Table S2-3 for statistical results and estimates.

Consistent with the pattern across all measurements, T_50_ increased with increasing elevation within families (Fig. 2; Supplementary Data Table S2-3), and within species (Fig. 3; Supplementary Data Table S2 and S4). T_50_ for each family was significantly higher at higher elevations, except for Costaceae (Fig. 2; Supplementary Data Table S2-3). *See* Supplementary Data Fig. S2 for heat tolerance of Zingiberales among all families. For 17 out of the 29 species that occur in more than two sites, T_50_ also increased with elevation (Fig. 3; Supplementary Data Table S2 and S4; R^2^ = 0.81, F = 20.13, P < 2.2E-16). For the rest of the species, T_50_ stayed the same or decrease with an increase of elevation (Fig. 3).

**Figure 3.**
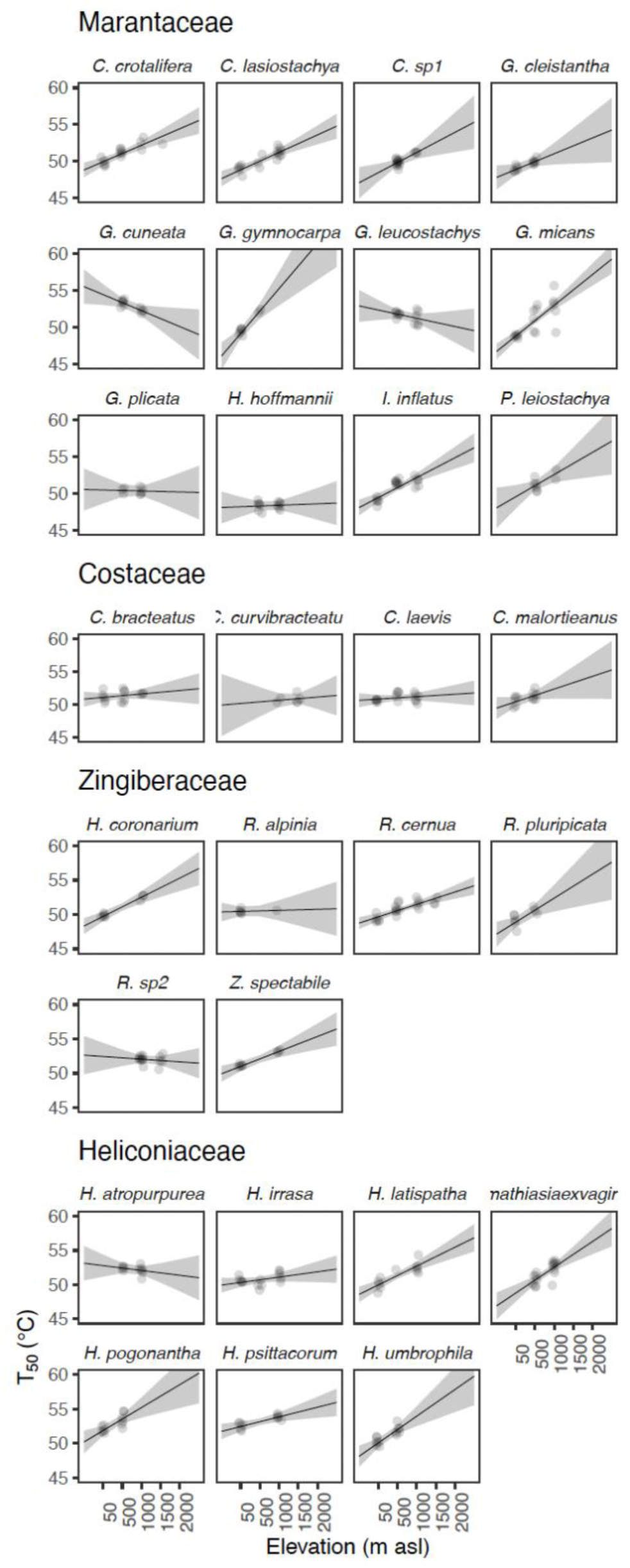
Relationship of heat tolerance between species with broad distributions and elevation at the Barva transect. Each dot represents T_50_ of a single individual, lines indicate linear regression from the model, and shaded areas indicate 95% confidence intervals of the predictions. *See* Supplementary Data Table S2 and S4 for statistical results and estimates.

The species with the highest heat tolerance was Heliconia psittacorum, found in the low premontane rainforest (1000 m asl) and in open areas (T_50_ = 53.9 ± 0.4°C, mean ± SD). This is an exotic species native to South America and is frequently used as an ornamental in gardens surrounding our study area. The species with the lowest T_50_ was Goeppertia venusta, a rare species found only in the shaded understory of the wet forest (50 m asl; T_50_ = 48.1 ± 0.2°C). We investigated whether species inhabiting sun-exposed or shaded environments displayed differential heat tolerance across each elevation. Our findings suggest that among Zingiberales, T_50_ of leaves of species found in full sunlight did not differ significantly from that of species in shaded areas (Supplementary Data Fig. S3).

Thermal safety margins calculated from air temperature appear to increase with elevation (*See* Section S1, Fig. S5). While leaf temperature would be a more reliable indicator for estimating thermal safety margins (Pau et al. 2018; Perez and Feeley 2020; Tarvainen et al. 2022), air temperature likely provides a reasonable approximation for our shade measured species. Given the accelerated rate of warming in tropical montane ecosystems, understanding the thermal safety margins of these species is valuable for predicting their future persistence.

## DISCUSSION

Our results indicate that leaf heat tolerance (as recorded by T_50_) changes among populations of Zingiberales along a tropical elevational gradient (Fig. 1). The general trend of T_50_ increasing with elevation is altered at the highest elevation of 2000 m with a single species, H. lankesterii. The low T_50_ at this elevation might reflect a chance event resulting from having only a single species here, as T_50_ varied widely within the community at all elevations. It might also reflect a true pattern of T_50_ peaking at mid-elevation, and the only Zingiberales species at 2000 m asl exhibiting significantly down-regulated heat tolerance. H. lankesterii is a 3 m tall understory plant restricted to the highest elevations of the transect. Compared to H. latispatha, which is distributed from Mexico to Perú, H. lankesterii has a more limited range confined to the highlands of Costa Rica and Panamá (Hammel et al. 2004).

We expected heat tolerance to be greater at lower elevations, where air temperatures are higher than at high-elevation sites. However, our results showed that heat tolerance for these herbaceous species increases with elevation, while air temperatures decrease. This trend was observed across plant communities and families, and within most species with broad distributions along the elevational gradient. In contrast, previous studies using the same techniques reported a reduction of heat tolerance of tropical tree species with increasing elevation, and heat tolerance to be associated with the maximum air temperature (Feeley et al. 2020; Slot et al. 2021; Tarvainen et al. 2022).

Besides temperature, many other factors also change with elevation, such as solar radiation, humidity, wind speed, CO_2_ partial pressure or rainfall. Solar radiation increases with increasing elevation, particularly the short-wave (UV-A and UV-B) component (Cavelier 1996; Smith and Geller 1979). Tropical mountains receive the highest natural levels of UV-B radiation on the planet, which is known to damage PSII (Bilger, Johnsen, and Schreiber 2001; Sicora, Máté, and Vass 2003; Ziska, Teramura, and Sullivan 1992; 1992), and adaptation to high UV radiation may contribute to an increased capacity to tolerate heat at higher elevations (Sullivan, Teramura, and Ziska 1992; Ziska, Teramura, and Sullivan 1992).

A recent study recorded T_50_ values of 45.4 - 53.9°C for a community of non-woody plant species at 3000 m elevation in tropical alpine highlands in the Andes (Leon-Garcia and Lasso 2019). These values are close to or higher than those reported in lowland tropical tree species (Feeley et al. 2020; Tiwari et al. 2021). High elevations such as the Andes experience extreme diurnal temperature shifts (Leon-Garcia and Lasso 2019), as well as high levels of potentially harmful UV radiation. In our study, we observed large changes in diurnal temperature at all the highest elevations (Supplementary Data Fig. S4 and S5). Adaptations to one type of physiological stress might increase protection against other physiological stressors (Levitt 1951). As such, the decreasing minimum temperature may have counterintuitively contributed to higher heat tolerance at higher elevations observed in the current study. Environmental stressors such as heat, cold, and UV radiation can lead to formation of reactive oxygen species (ROS) which are highly reactive and toxic, mainly produced in PSII and PSI, which ultimately results in cell damage (Gill and Tuteja 2010). In turn, ROS production during stress periods has been suggested to play a central role in stress perception and protection, even interlinked with temperature tolerance and acclimation (Suzuki and Mittler 2006). T_50_ is a measurement of the physical state of the chloroplast membrane. It is associated with the PSII reaction center and its associated light harvesting component, all physical structures that are affected by environmental variables. Thus, while T_50_ is a good measure of heat tolerance, it may also reflect changes in the physical composition of the membrane and PSII influenced by other environmental or genetic factors. In the current study, adaptation to increased UV exposure along the Barva gradient could have led to higher heat tolerance as a ‘byproduct’, at least in the sun-exposed species. Still, more research is needed to determine if variation in solar radiation is a factor determining leaf temperature at the Barva transect, if UV-B radiation increases with elevation, and its link to higher tolerance of PSII in Zingiberales.

Results from previous studies support the hypothesis that heat tolerance should match the extreme temperatures that plants experience daily (Feeley et al. 2020; Perez et al. 2021). However, higher elevations experience a wide range of temperatures, with summer-like conditions during the day and winter-like conditions during the night (Ghalambor 2006; Mani 2013). This daily temperature fluctuation might be the driver for such a high heat tolerance in herbaceous plants, contrary to the general expectation of the lowlands having higher heat tolerances because of their high mean annual temperatures. The heat tolerance among herbaceous plants such as Zingiberales and other non-woody species (Leon-Garcia and Lasso 2019) seems higher than for woody species at high elevations. Additionally, it has been shown that low temperatures result in photoinhibition of PSII, especially when F_v_/F_m_ is measured in herbaceous plants growing in full sun (Germino and Smith 2000). Lower temperatures at higher elevations could potentially lead to cold-induced photoinhibition, as it has been shown in mango cultivars, which have been shown to experience cold-induced photoinhibition at 10°C and lower F_v_/F_m_ values over time (Sukhvibul et al. 2000). The minimum temperatures at higher elevation sites might be sufficiently low to induce photoinhibition in principle. However, our control leaves for F_v_/F_m_ values (> 0.7) suggest minimal photoinhibition, which may be an adaptation to cold or variable temperatures. This adaptation could potentially provide some benefit for heat tolerance as well. Additional studies are needed to elucidate these differences among different growth forms in the same location, and the role of maximum and minimum temperature in controlling heat tolerance.

Additional factors that may influence heat tolerance in Zingiberales include those that decouple leaf temperature from air temperature (Feeley et al. 2020; Michaletz et al. 2015). For example, leaf size and shape affect leaf temperature. Larger and wider leaves can reach higher leaf temperatures than smaller and thinner leaves (Leigh et al. 2017; I. J. Wright et al. 2017). Zingiberales species have very different leaf sizes. For example, leaves of Heliconia pogonantha can be 3 m long. In contrast, Goeppertia micans leaves are at most 15 cm long (Hammel et al. 2004). Many Zingiberales species also display leaf behaviors that prevent higher temperatures and avoid heat stress. For example, marantaceas can rotate their leaves upright by changing the turgor of the pulvinus (Herbert and Larsen 1985), thereby reducing radiation loads. Plants in the genus Musa and Heliconia frequently tear, resulting in the large leaf blades that are common in these families being subdivided into smaller sections (Taylor and Sexton 1972), significantly reducing their boundary layer. These leaf adaptations could explain some of the differences in heat tolerance among species and families but not the general pattern of increasing heat tolerance with elevation among populations of the same species.

Higher temperatures are assumed to severely impact the tropics, especially the lowlands (Stocker 2014; S. J. Wright, Muller-Landau, and Schipper 2009). Lowland plants may be at higher risk because they are already experiencing high temperatures and humidity, which is expected to reach deleterious levels in the next century (Doughty et al. 2023; Scheffers et al. 2014). Rising vapor pressure deficit (Barkhordarian et al. 2019) could increase plant stressors, especially for lowlands species. Vapor pressure deficit increases exponentially with temperature, independent of the rainfall (de Boer et al. 2011). If there is an increase in vapor pressure deficit it can cause stomata to close and reduce the potential of carbon assimilation (Grossiord et al. 2020; Slot et al. 2024; Slot and Winter 2017). Then, in a cascading effect, closed stomata may lead to an increase in leaf temperature because latent heat loss is reduced because of the decrease in transpiration. In contrast, high-elevation regions are expected to experience more rapid temperature increases due to climate change than low-elevation areas (Pepin, N et al. 2015). If this is the case, high heat tolerance of species in higher elevations could be a beneficial trait to withstand higher air temperatures.

## CONCLUSIONS

Heat tolerance of Zingiberales increases with elevation. Our estimates of T_50_ integrate an unknown combination of physiological adaptation and plasticity. This variation within species and among populations allows plants to persist in heterogeneous and stochastic environments (Ahrens et al. 2021). Although an increase in heat tolerance is not obviously advantageous for species migrating to higher elevations, it is possible that it can facilitate colonization when exposed to extreme temperatures in novel environments. However, if plants already have high heat tolerance, which can also be plastic and increase over time, species may not require colonizing novel environments. Heat tolerance is the result of many complex physiological and morphological traits (Lortie et al. 2004; Mitra and Bhatia 2008). Our results illustrate that heat tolerance may reflect a response to different environmental stressors not accounted for in this study, affecting the physical structures within the photosynthetic apparatus and cause increases in apparent heat tolerance.

## SUPPLEMENTAL MATERIAL

Supplementary data are available online at XX and consist of the following. **Section S1:** Thermal Safety Margins of Zingiberales along the Barva transect. **Table S1.** List of species, abbreviations of the species respectively, elevations at which each species was collected, and average heat tolerance for Zingiberales species included in the study. **Table S2.** Results of analyses of the ANOVA for each of the models. **Table S3.** Estimated marginal means of linear trends. **Table S4.** Estimates to show differences of T_50_ along the gradient within species. **Figure S1.** Experiment set up to determine heat tolerance (T_50_). (a) Water baths with submerged leaf tissue placed in plastic bags. (b) Leaf disks in miracloth fabric and then placed in bags ready for submersion. (c) After temperature treatment in the water baths, leaf tissue is placed in a petri dish for 24h. **Figure S2.** Heat tolerance of Zingiberales families along the Barva transect. Heat tolerance (T_50_) was estimated as the temperature at which PSII quantum yield (fluorescence) decreases 50%. Plant families in phylogenetic order: Musaceae (MU), Heliconiaceae (HE), Cannaceae (CA), Marantaceae (MA), Costaceae (CO), and Zingiberaceae (ZI). Error bars indicate standard deviation, boxes indicate standard error, and the central line represents average heat tolerance for each family. Dots represent the mean of each species at the given family. **Figure S3.** Heat tolerance (T_50_) of Zingiberales individuals as a function of elevation at the Barva transect. Dots represent heat tolerance of individuals from each species at the given elevation site. Grey and yellow dots indicate if the species were collected from a shaded environment or a fully sun environment, respectively (see Table S1 for complete information on species environment). **Figure S4.** Maximum and minimum daily temperature (blue and red dots, respectively) for each elevation by month (Clark & Clark 2015) at the Barva transect, Costa Rica. **Figure S5.** (a) Heat tolerances of Zingiberales along the Barva transect (black lines at the top of the figure), and maximum and minimum daily temperature (blue and red bars and dots near the bottom of the figure) for each elevation. Error bars indicate standard deviation, and the central horizontal line represents average heat tolerance for plant communities at each elevation. For the temperature data in the bottom, the central dark square represents the average for each maximum (red) and minimum (blue) daily temperature each elevation, and each dot represents the maximum and minimum daily temperature. (b) Thermal Safety Margins for each elevation at the Barva transect. Each dot represents TSM individual plants at each elevation. The line indicates linear regression from the model. Note that at the highest elevation only one species is present.

## FUNDING

This work was supported by funds from the Glaxo Centro America Research Fellowship [Fund 502] of the Organization for Tropical Studies, Heliconia Society International Award for Botanical and Horticultural Research Projects on the Zingiberales, Ronald Bamford Fund to the Department of Ecology and Evolutionary Biology, University of Connecticut, and The Explorers Club – Mamont Scholar Grant to G.G.H, grants from the National Science Foundation (Dimensions of Biodiversity grant #1737778, Organismal Responses to Climate Change #2222328) to CG-R.

## ACKNOWLEDGMENTS

We thank the staff of La Selva Biological Station Organization for Tropical Studies for logistical support, Orlando Vargas for guidance in species identification, Marcos Molina, Pablo Muñoz, and Constantino Hernández Cordero for valuable assistance in the field. We thank Carl Schlichting, Laura Bizzarri, and Erin Kuprewicz for their valuable comments on the manuscript. We thank the Sistema Nacional de Áreas de Conservación for permission to conduct research in Costa Rica (SINAC-ACC-PI-R-044-2019).

## AUTHOR CONTRIBUTIONS

GGH and CGR conceived the ideas, GGH planned and designed the research; GGH collected the data; GGH analyzed the data with contributions from LdCG, GGH wrote the manuscript with substantial contributions from MS, CGR and JRS. All authors contributed critically to the drafts.

## OPEN RESEARCH STATEMENT

Data used in this paper are available at Zenodo repository: Hernández, G. G. (2024). Heat tolerance of tropical herbaceous plants increases with elevation [Data set]. Zenodo. https://doi.org/10.5281/zenodo.13892118. The code is available at: https://github.com/georgiahernandez/HeatTolerance_BarvaElevation

## CONFLICT OF INTEREST STATEMENT

The authors declare no conflict of interest.

## Section S1

### Thermal Safety Margins of Zingiberales along the Barva transect

To determine differences in vulnerabilities to global warming among individuals at each elevation, we estimated the thermal safety margins (TSM) of all Zingiberales species along the Barva transect (Fig. S5). TSM is the difference between maximum air temperature and T_50_. Individuals from the lowlands tended to have lower thermal safety margins than individuals from higher elevations (TSM = 14.82 + 0.008 × Elevation, F = 1512, R2 = 0.75, P < 2.2E-16).

Heat tolerance and the temperature range at each site of the Barva gradient is shown in Fig. S5A. In general, thermal limits of all Zingiberales species were at least 12°C above their respective air temperature (Fig. S5B). Species found in the premontane rainforest (HPR, 1500 m asl) displayed the highest thermal safety margins compared to other life zones at their specific elevations (Fig. 4B). We recorded the following TSM mean and standard deviation for all individuals at each elevation: 50 m asl TSM = 14.1 ± 1.3°C (N = 200), 500 m asl TSM = 21.4 ± 1.4°C (N = 139), 1000 m asl TSM = 21.9 ± 1.4°C (N = 130), 1500 m asl TSM = 27.6 ± 1.2°C (N = 31), and 2000 m asl TSM = 23.5 ± 0.6°C (N = 6).

Our estimates of thermal safety margins are very conservative, as they are based on air temperature. Recent investigations in tropical trees have advocated for the incorporation of leaf temperature, rather than ambient or environmental temperature, as the key determinant of plant functioning (Pau et al. 2018; Perez and Feeley 2020; Tarvainen et al. 2022). This shift is substantiated by the observation that air temperature data obtained from weather stations exhibits limited congruence with leaf temperature, as they do not necessarily correlate closely (Doughty et al. 2023; Grace, Berninger, and Nagy 2002; Pau et al. 2018; Perez and Feeley 2020; Tserej and Feeley 2021). Furthermore, empirical evidence indicates that thermal tolerances are predominantly associated with leaf temperature, as opposed to ambient air temperature (Perez and Feeley 2020). In fact, leaf temperature could be greatly decoupled from air temperature by plant thermoregulatory processes and morphology, as previously mentioned.

---

**Table S1.**
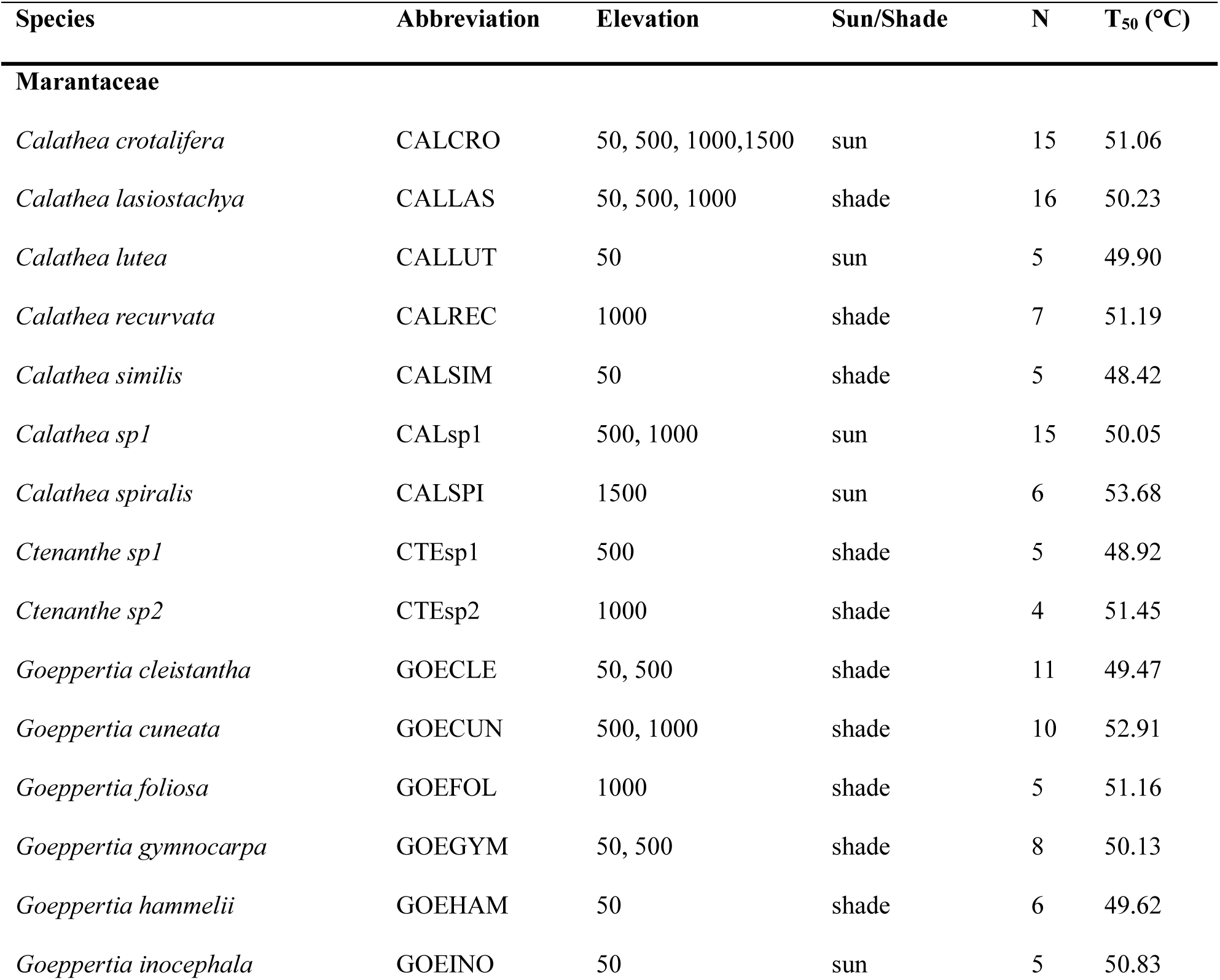
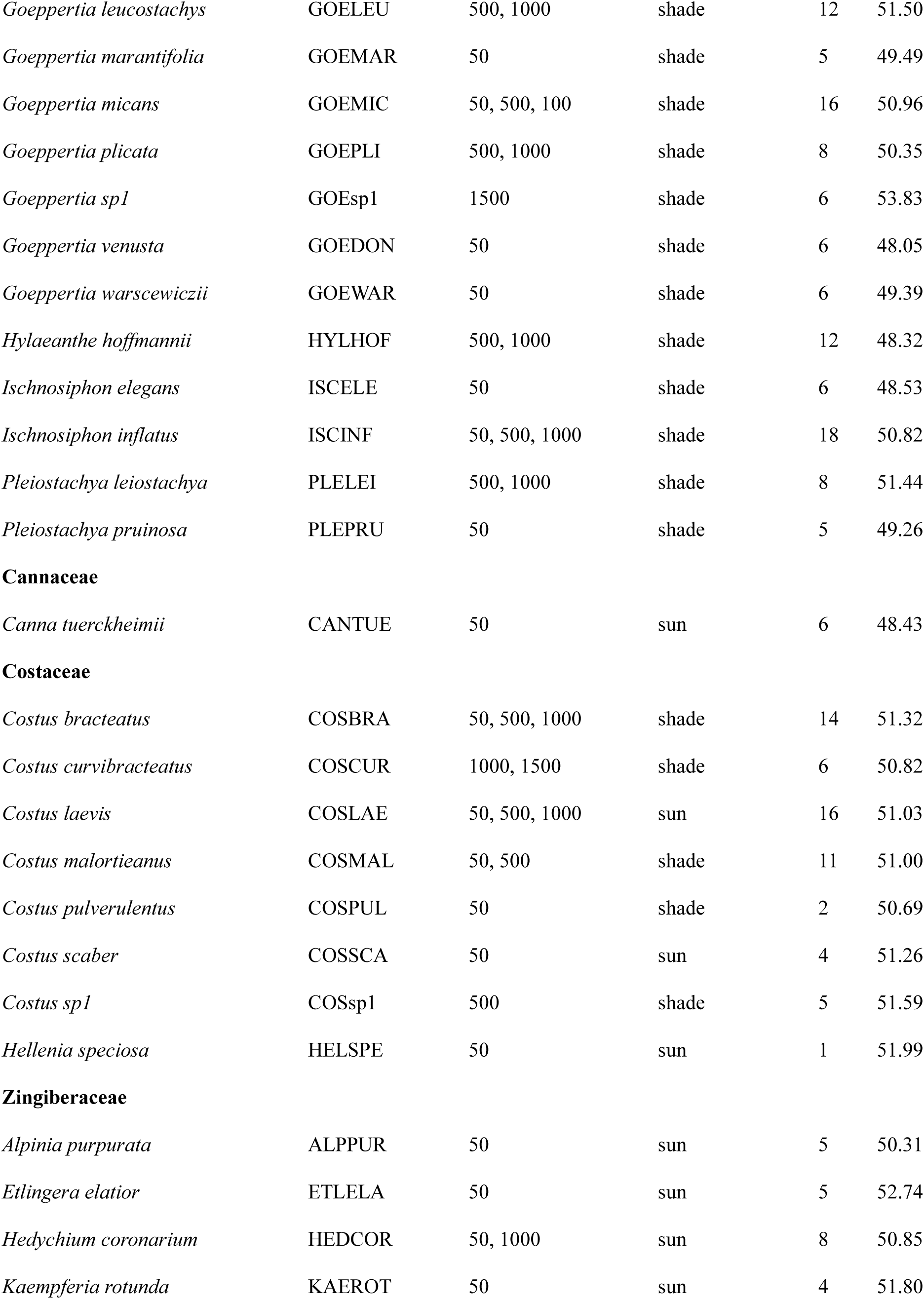
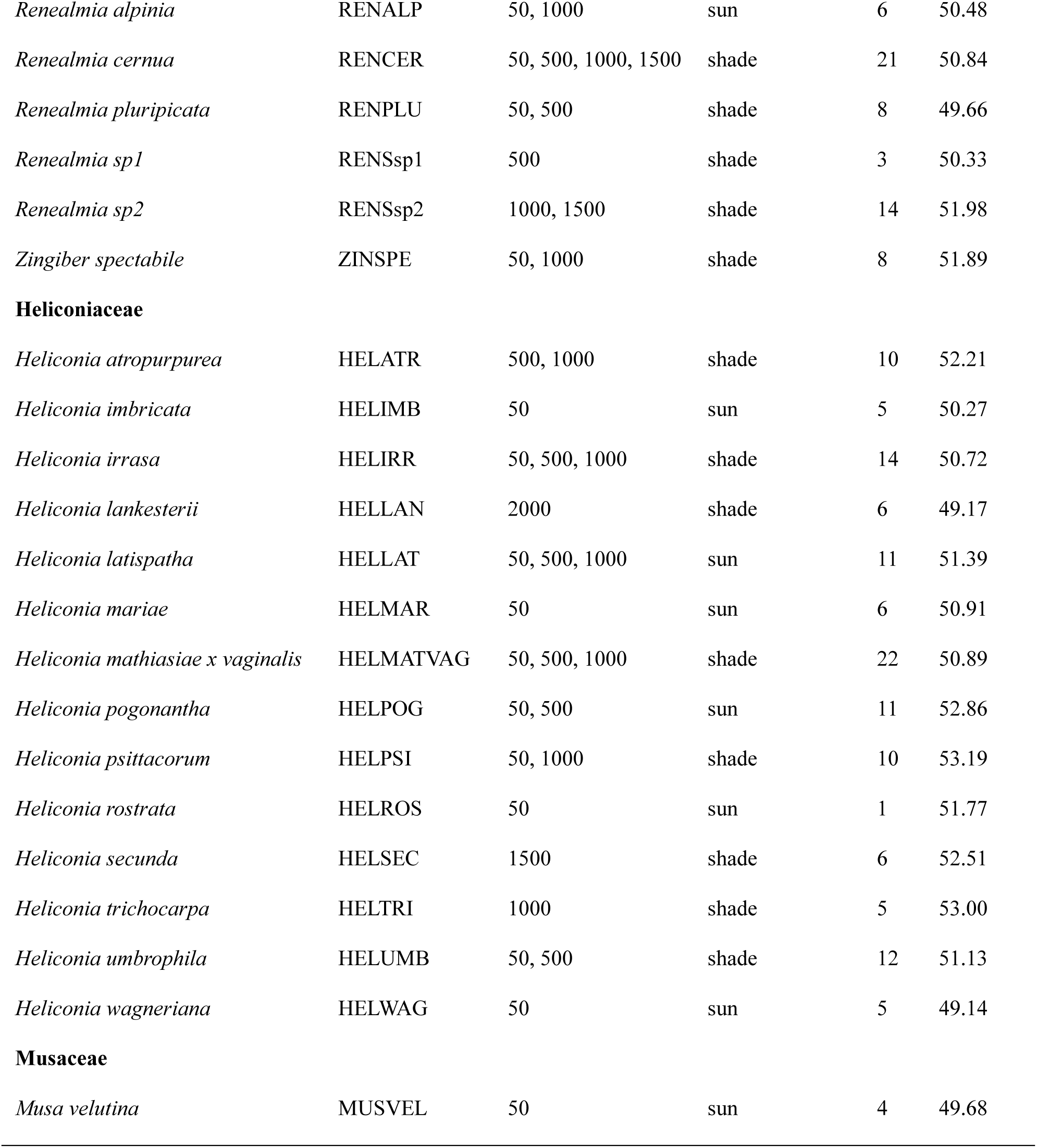
List of species, abbreviations of the species respectively, elevations at which each species was collected, and average heat tolerance for Zingiberales species included in the study. Sample size (N) is the number of individuals sampled from each species at all elevations. Sun/Shade is the environment where species were sampled along the elevational gradient. Sun conditions include recently abandoned pastures, gardens and any open area with full sun conditions. Shade conditions include any percent of canopy coverage and may have varied along the transect within the same species. T_50_ estimates represent the average for each species at all elevations.

**Table S2.**
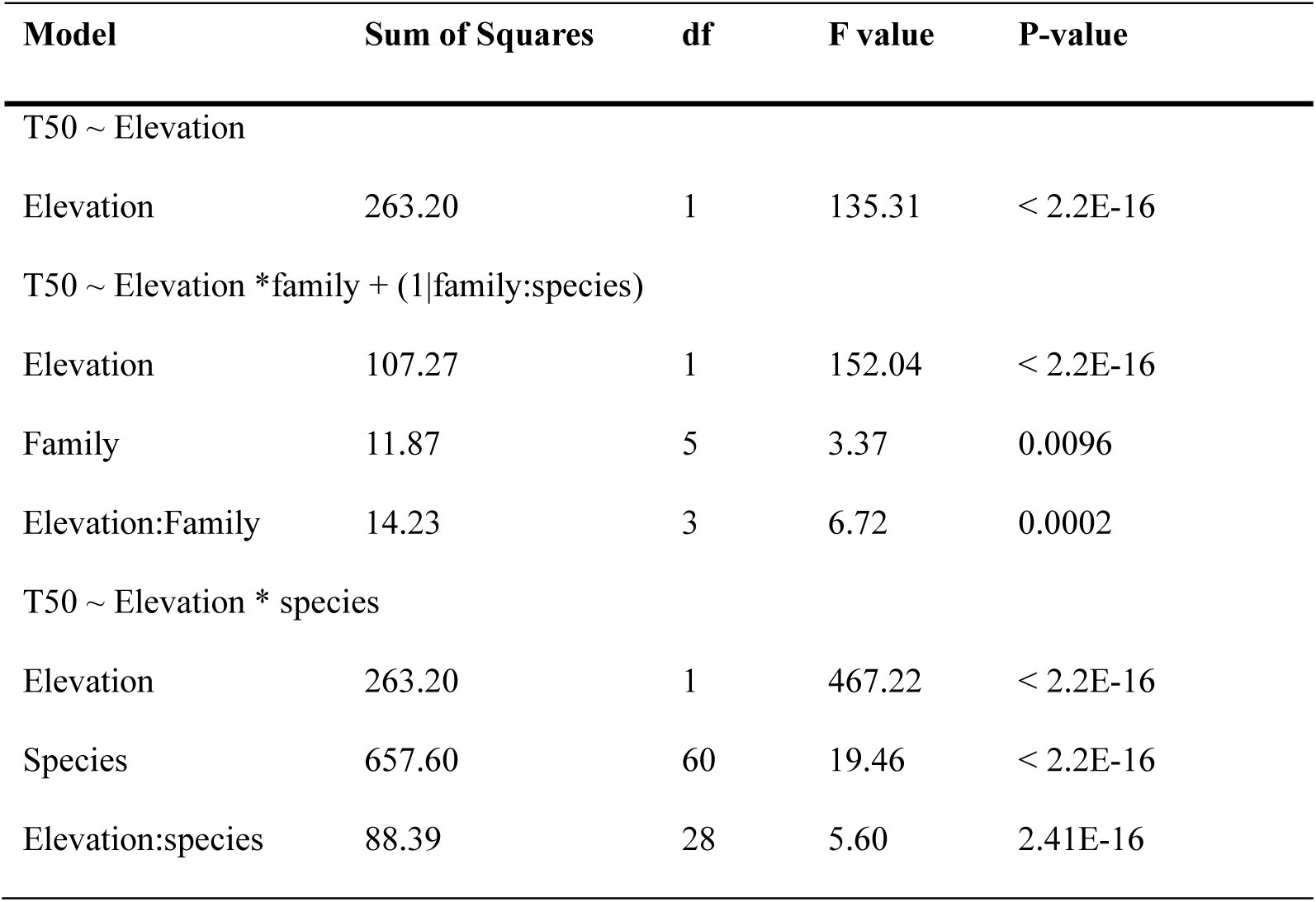
Results of analyses of the ANOVA for each of the models.

**Table S3.**
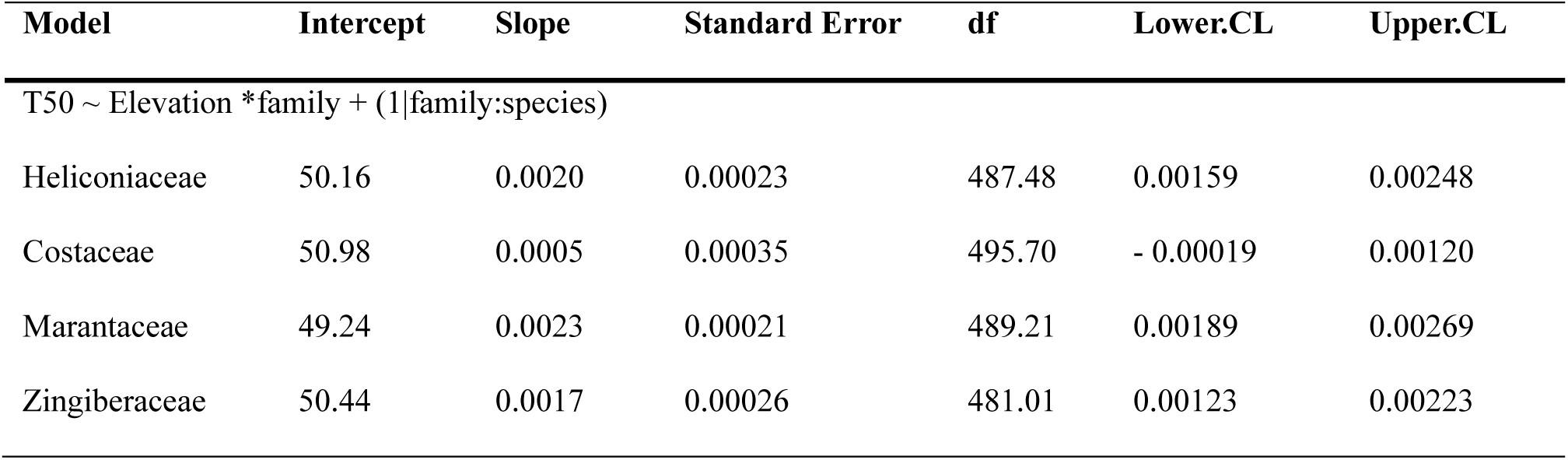
Estimated marginal means of linear trends.

**Table S4.**
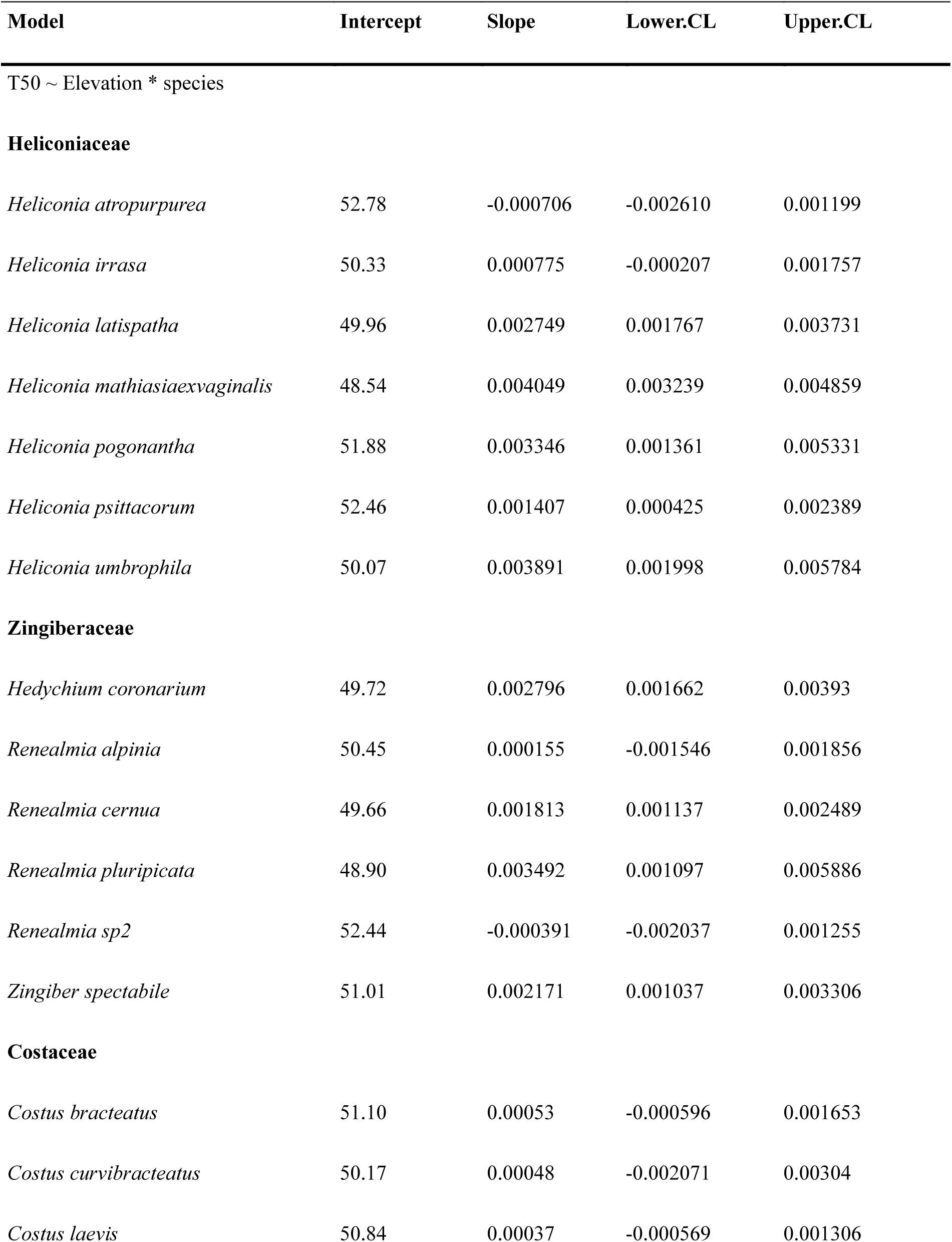
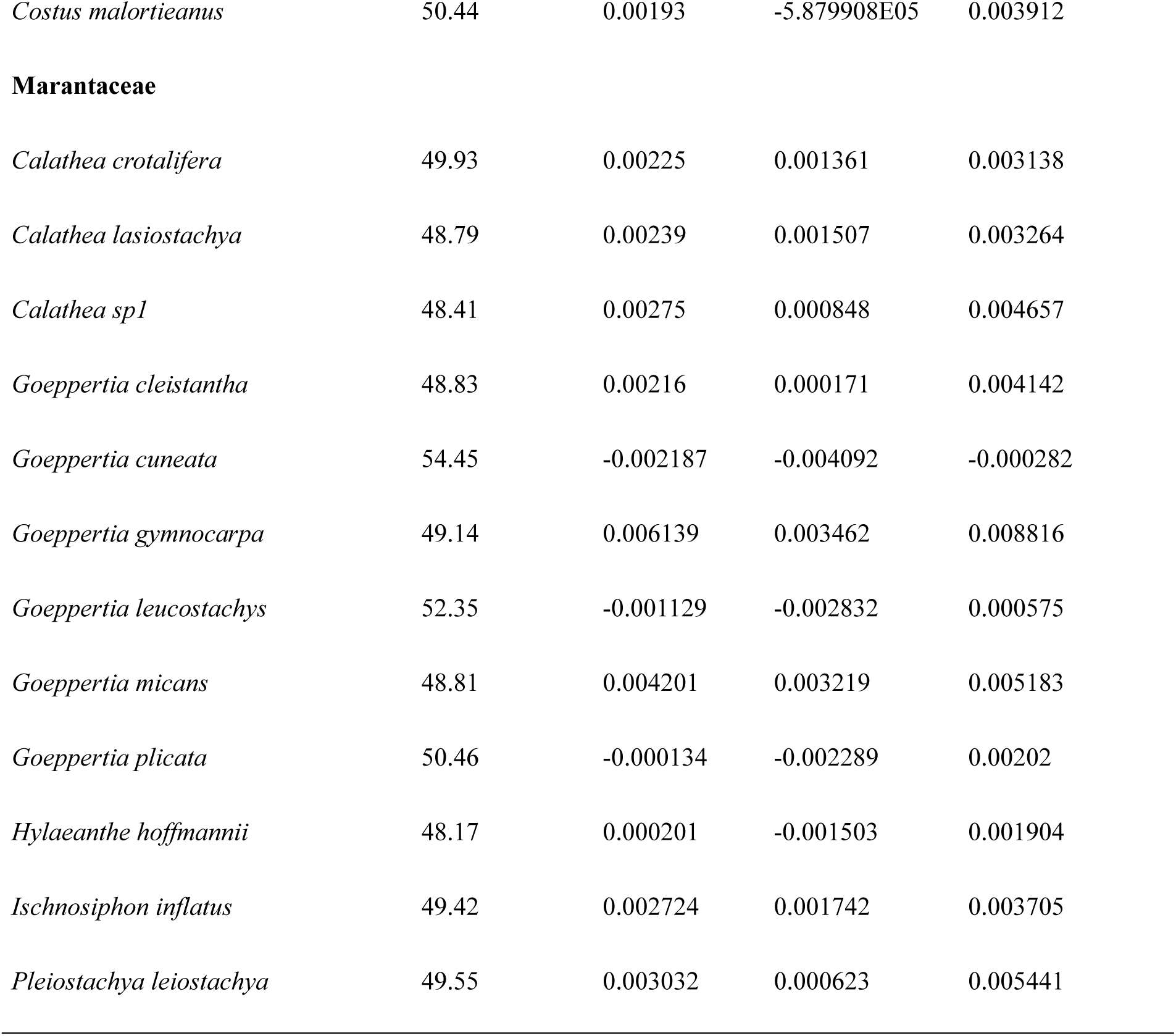
Estimates to show differences of T_50_ along the gradient within species.

**Figure S1.**
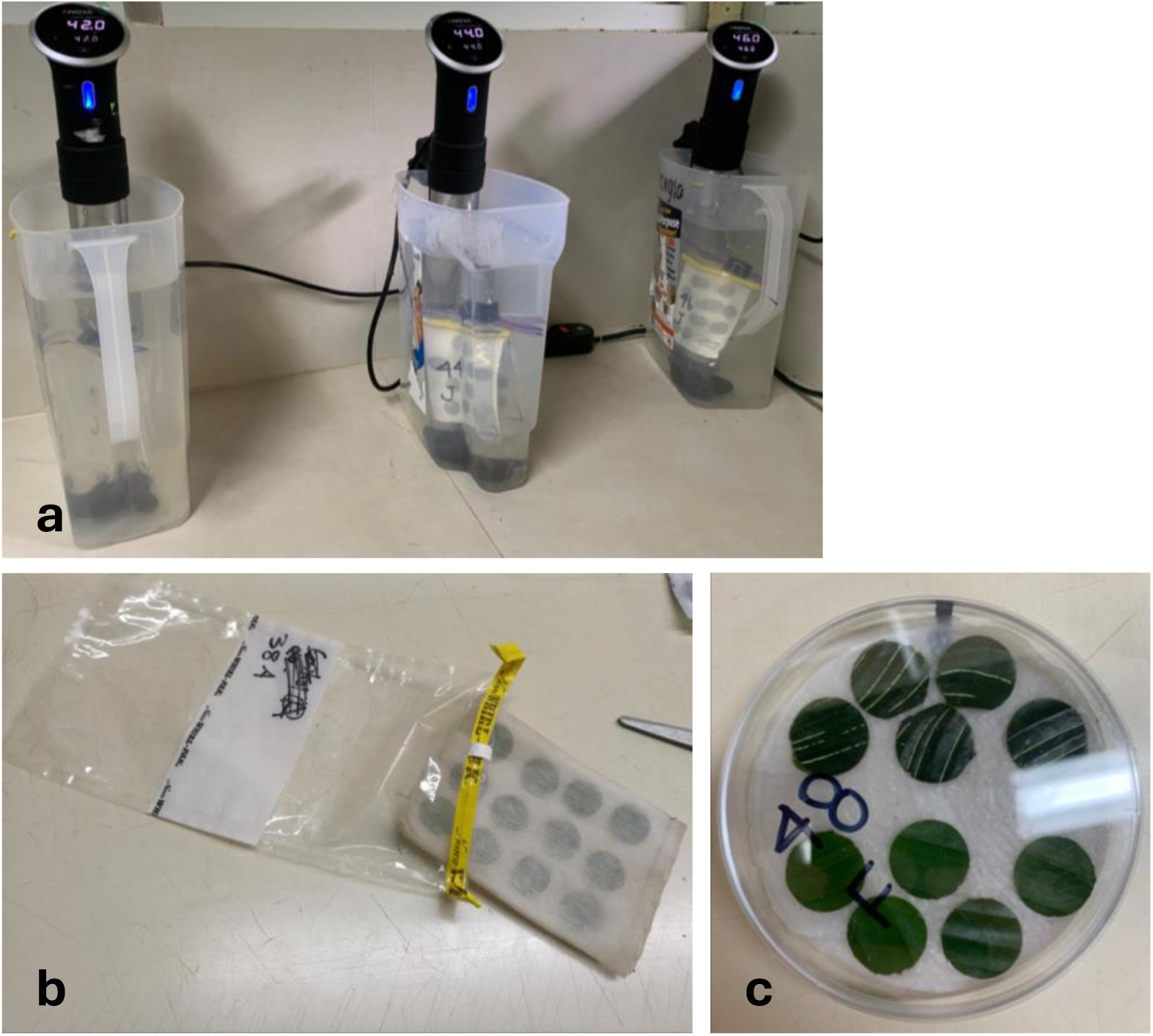
Experiment set up to determine heat tolerance (T_50_). **(a)** Water baths with submerged leaf tissue placed in plastic bags. **(b)** Leaf disks in miracloth fabric and then placed in bags ready for submersion. **(c)** After temperature treatment in the water baths, leaf tissue is placed in a petri dish for 24h.

**Figure S2.**
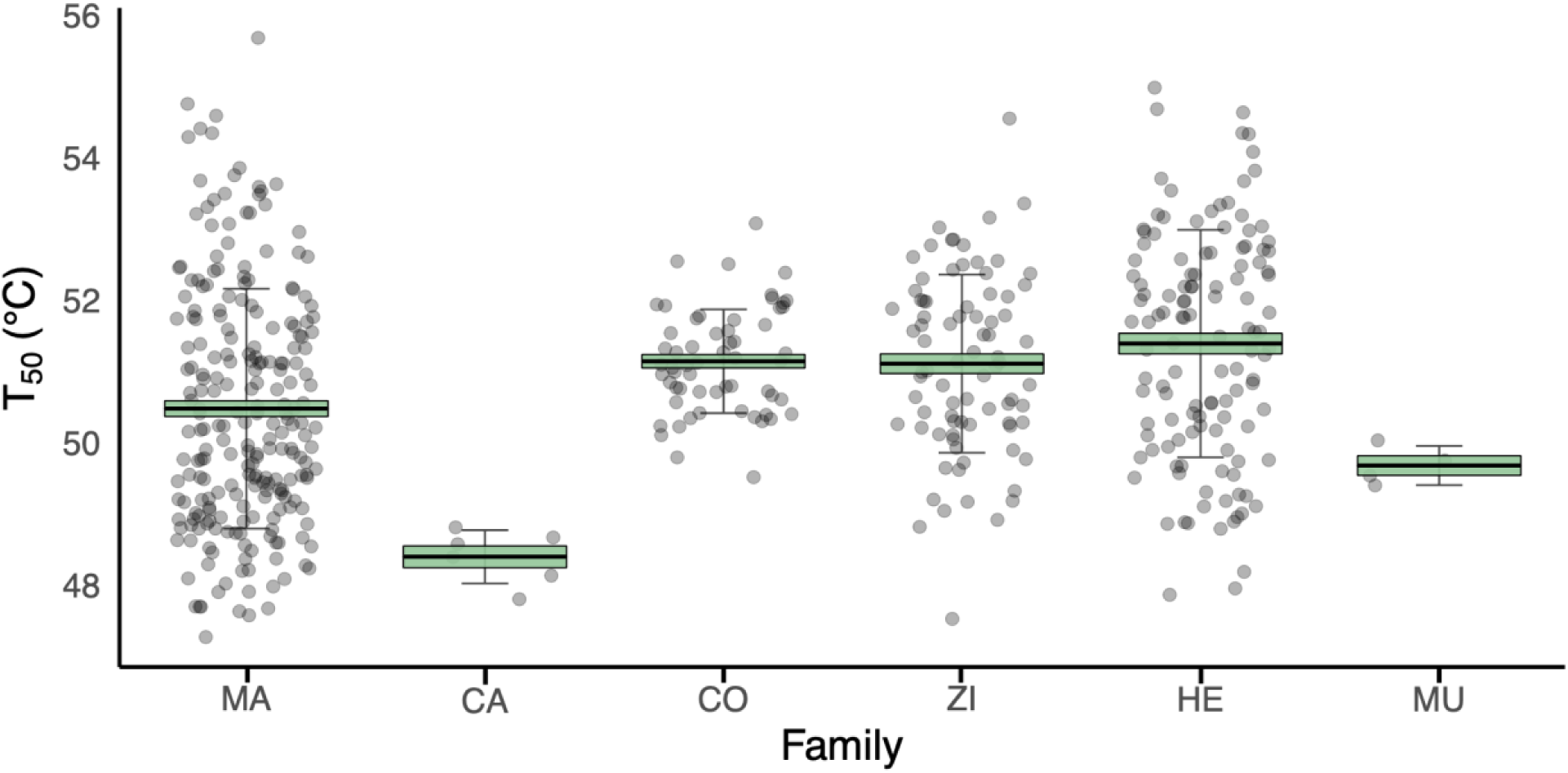
Heat tolerance of Zingiberales families along the Barva transect. Heat tolerance (T_50_) was estimated as the temperature at which PSII quantum yield (fluorescence) decreases 50%. Plant families in phylogenetic order: Musaceae (MU), Heliconiaceae (HE), Cannaceae (CA), Marantaceae (MA), Costaceae (CO), and Zingiberaceae (ZI). Error bars indicate standard deviation, boxes indicate standard error, and the central line represents average heat tolerance for each family. Dots represent the mean of each species at the given family.

**Figure S3.**
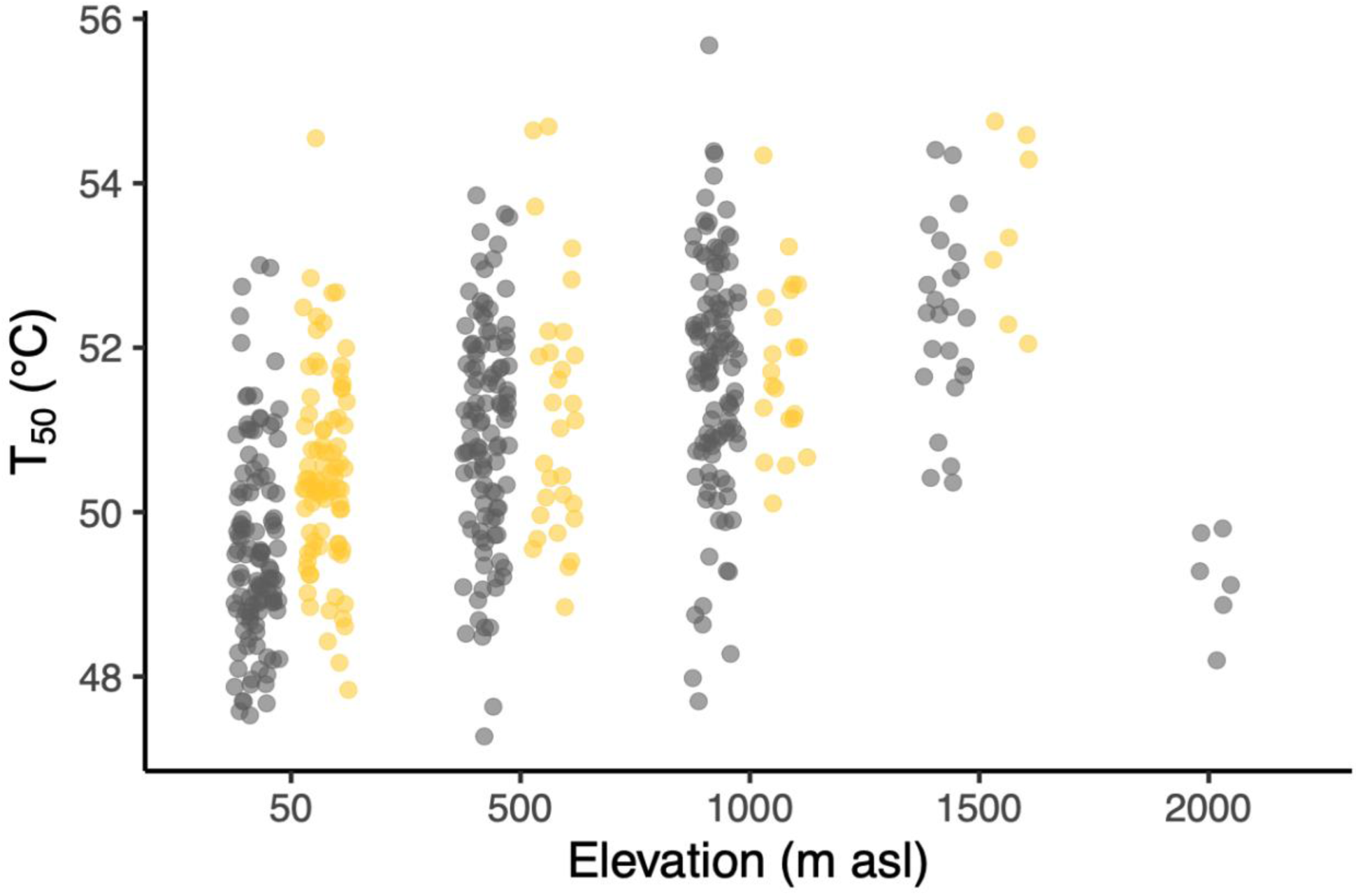
Heat tolerance (T_50_) of Zingiberales individuals as a function of elevation at the Barva transect. Dots represent heat tolerance of individuals from each species at the given elevation site. Grey and yellow dots indicate if the species were collected from a shaded environment or a fully sun environment, respectively (see Table S1 for complete information on species environment).

**Figure S4.**
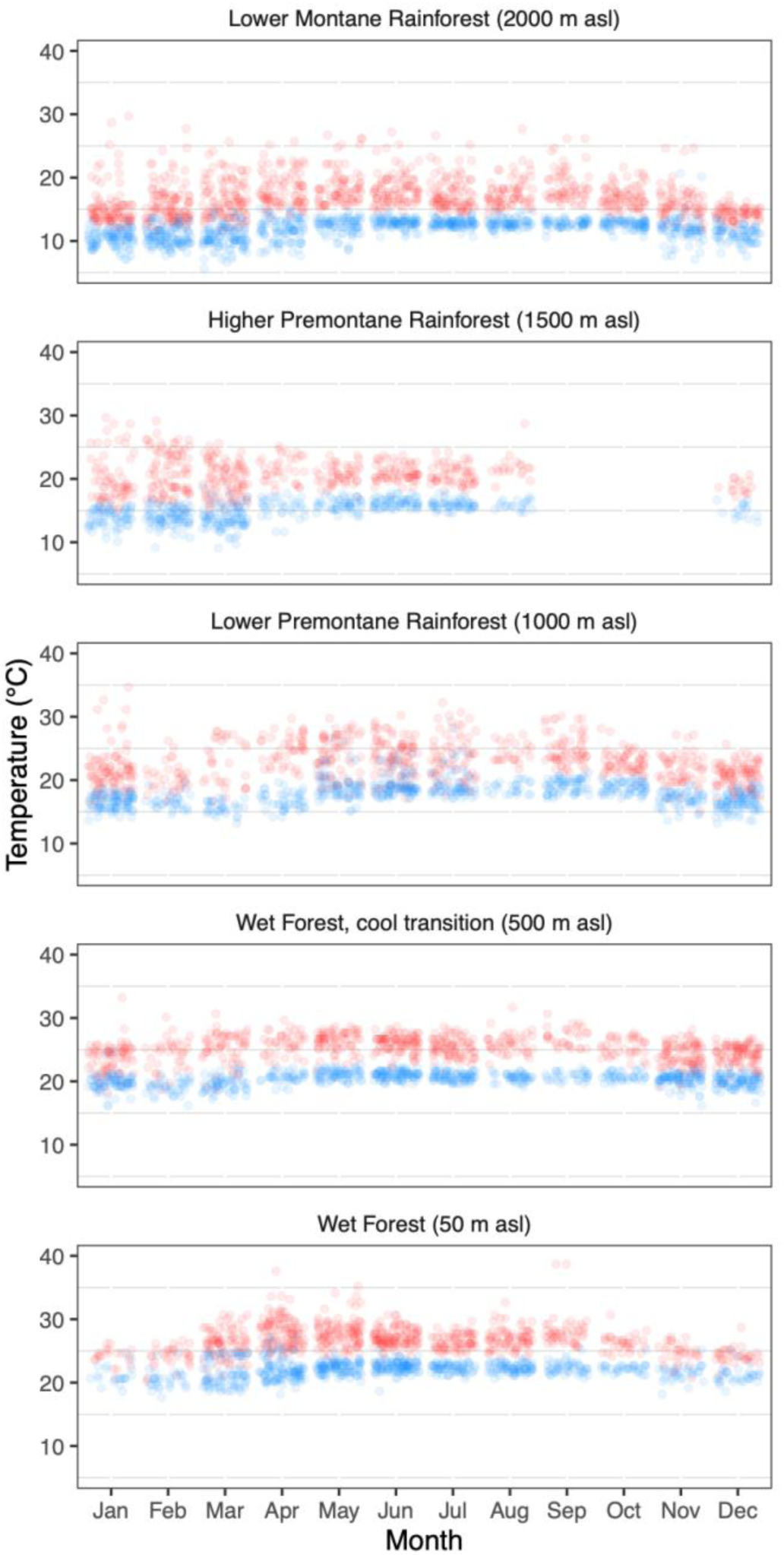
Maximum and minimum daily temperature (blue and red dots, respectively) for each elevation by month (Clark & Clark 2015) at the Barva transect, Costa Rica.

**Figure S5.**
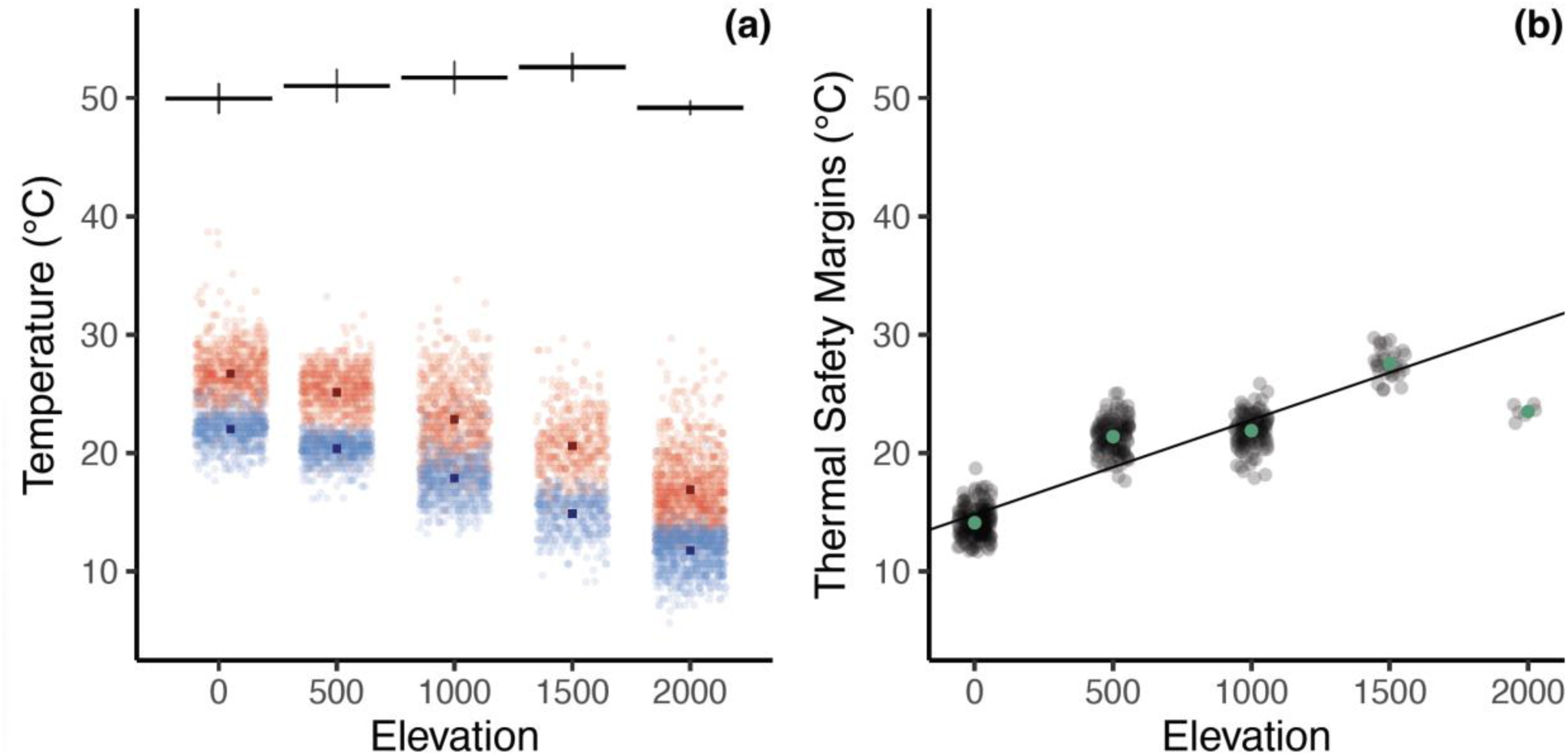
**(a)** Heat tolerances of Zingiberales along the Barva transect (black lines at the top of the figure), and maximum and minimum daily temperature (blue and red bars and dots near the bottom of the figure) for each elevation. Error bars indicate standard deviation, and the central horizontal line represents average heat tolerance for plant communities at each elevation. For the temperature data in the bottom, the central dark square represents the average for each maximum (red) and minimum (blue) daily temperature each elevation, and each dot represents the maximum and minimum daily temperature. **(b)** Thermal Safety Margins for each elevation at the Barva transect. Each dot represents TSM individual plants at each elevation. The line indicates linear regression from the model. Note that at the highest elevation only one species is present. *See* Supplementary Data Section S1 for the TSM methods and statistical results.

## Notes

### Competing Interest Statement

The authors have declared no competing interest.

