## Supplementary Materials for "Heat tolerance of tropical herbaceous plants increases with elevation"

**Table S1.** List of species, abbreviations of the species respectively, elevations at which each species was collected, and average heat tolerance for Zingiberales species included in the study. Sample size (N) is the number of individuals sampled from each species at all elevations. Sun/Shade is the environment where species were sampled along the elevational gradient. Sun conditions include recently abandoned pastures, gardens and any open area with full sun conditions. Shade conditions include any percent of canopy coverage and may have varied along the transect within the same species. $T_{50}$ estimates represent the average for each species at all elevations.

| Species | Abbreviation | Elevation | Sun/Shade | N | $T_{50}$ (°C) |
|---|---|---|---|---|---|
| **Marantaceae** | | | | | |
| *Calathea crotalifera* | CALCRO | 50, 500, 1000,1500 | sun | 15 | 51.06 |
| *Calathea lasiostachya* | CALLAS | 50, 500, 1000 | shade | 16 | 50.23 |
| *Calathea lutea* | CALLUT | 50 | sun | 5 | 49.90 |
| *Calathea recurvata* | CALREC | 1000 | shade | 7 | 51.19 |
| *Calathea similis* | CALSIM | 50 | shade | 5 | 48.42 |
| *Calathea sp1* | CALsp1 | 500, 1000 | sun | 15 | 50.05 |
| *Calathea spiralis* | CALSPI | 1500 | sun | 6 | 53.68 |
| *Ctenanthe sp1* | CTEsp1 | 500 | shade | 5 | 48.92 |
| *Ctenanthe sp2* | CTEsp2 | 1000 | shade | 4 | 51.45 |
| *Goeppertia cleistantha* | GOECLE | 50, 500 | shade | 11 | 49.47 |
| *Goeppertia cuneata* | GOECUN | 500, 1000 | shade | 10 | 52.91 |
| *Goeppertia foliosa* | GOEFOL | 1000 | shade | 5 | 51.16 |
| *Goeppertia gymnocarpa* | GOEGYM | 50, 500 | shade | 8 | 50.13 |
| *Goeppertia hammelii* | GOEHAM | 50 | shade | 6 | 49.62 |
| *Goeppertia inocephala* | GOEINO | 50 | sun | 5 | 50.83 |

| | | | | | |
|---|---|---|---|---|---|
| *Goeppertia leucostachys* | GOELEU | 500, 1000 | shade | 12 | 51.50 |
| *Goeppertia marantifolia* | GOEMAR | 50 | shade | 5 | 49.49 |
| *Goeppertia micans* | GOEMIC | 50, 500, 100 | shade | 16 | 50.96 |
| *Goeppertia plicata* | GOEPLI | 500, 1000 | shade | 8 | 50.35 |
| *Goeppertia sp1* | GOEsp1 | 1500 | shade | 6 | 53.83 |
| *Goeppertia venusta* | GOEDON | 50 | shade | 6 | 48.05 |
| *Goeppertia warscewiczii* | GOEWAR | 50 | shade | 6 | 49.39 |
| *Hylaeanthe hoffmannii* | HYLHOF | 500, 1000 | shade | 12 | 48.32 |
| *Ischnosiphon elegans* | ISCELE | 50 | shade | 6 | 48.53 |
| *Ischnosiphon inflatus* | ISCINF | 50, 500, 1000 | shade | 18 | 50.82 |
| *Pleiostachya leiostachya* | PLELEI | 500, 1000 | shade | 8 | 51.44 |
| *Pleiostachya pruinosa* | PLEPRU | 50 | shade | 5 | 49.26 |
| **Cannaceae** | | | | | |
| *Canna tuerckheimii* | CANTUE | 50 | sun | 6 | 48.43 |
| **Costaceae** | | | | | |
| *Costus bracteatus* | COSBRA | 50, 500, 1000 | shade | 14 | 51.32 |
| *Costus curvibracteatus* | COSCUR | 1000, 1500 | shade | 6 | 50.82 |
| *Costus laevis* | COSLAE | 50, 500, 1000 | sun | 16 | 51.03 |
| *Costus malortieanus* | COSMAL | 50, 500 | shade | 11 | 51.00 |
| *Costus pulverulentus* | COSPUL | 50 | shade | 2 | 50.69 |
| *Costus scaber* | COSSCA | 50 | sun | 4 | 51.26 |
| *Costus sp1* | COSsp1 | 500 | shade | 5 | 51.59 |
| *Hellenia speciosa* | HELSPE | 50 | sun | 1 | 51.99 |
| **Zingiberaceae** | | | | | |
| *Alpinia purpurata* | ALPPUR | 50 | sun | 5 | 50.31 |
| *Etlingera elatior* | ETLELA | 50 | sun | 5 | 52.74 |
| *Hedychium coronarium* | HEDCOR | 50, 1000 | sun | 8 | 50.85 |
| *Kaempferia rotunda* | KAEROT | 50 | sun | 4 | 51.80 |

| | | | | | |
|---|---|---|---|---|---|
| *Renealmia alpinia* | RENALP | 50, 1000 | sun | 6 | 50.48 |
| *Renealmia cernua* | RENCER | 50, 500, 1000, 1500 | shade | 21 | 50.84 |
| *Renealmia pluripicata* | RENPLU | 50, 500 | shade | 8 | 49.66 |
| *Renealmia sp1* | RENSsp1 | 500 | shade | 3 | 50.33 |
| *Renealmia sp2* | RENSsp2 | 1000, 1500 | shade | 14 | 51.98 |
| *Zingiber spectabile* | ZINSPE | 50, 1000 | shade | 8 | 51.89 |
| **Heliconiaceae** | | | | | |
| *Heliconia atropurpurea* | HELATR | 500, 1000 | shade | 10 | 52.21 |
| *Heliconia imbricata* | HELIMB | 50 | sun | 5 | 50.27 |
| *Heliconia irrasa* | HELIRR | 50, 500, 1000 | shade | 14 | 50.72 |
| *Heliconia lankesterii* | HELLAN | 2000 | shade | 6 | 49.17 |
| *Heliconia latispatha* | HELLAT | 50, 500, 1000 | sun | 11 | 51.39 |
| *Heliconia mariae* | HELMAR | 50 | sun | 6 | 50.91 |
| *Heliconia mathiasiae x vaginalis* | HELMATVAG | 50, 500, 1000 | shade | 22 | 50.89 |
| *Heliconia pogonantha* | HELPOG | 50, 500 | sun | 11 | 52.86 |
| *Heliconia psittacorum* | HELPSI | 50, 1000 | shade | 10 | 53.19 |
| *Heliconia rostrata* | HELROS | 50 | sun | 1 | 51.77 |
| *Heliconia secunda* | HELSEC | 1500 | shade | 6 | 52.51 |
| *Heliconia trichocarpa* | HELTRI | 1000 | shade | 5 | 53.00 |
| *Heliconia umbrophila* | HELUMB | 50, 500 | shade | 12 | 51.13 |
| *Heliconia wagneriana* | HELWAG | 50 | sun | 5 | 49.14 |
| **Musaceae** | | | | | |
| *Musa velutina* | MUSVEL | 50 | sun | 4 | 49.68 |

**Table S2.** Results of analyses of the ANOVA for each of the models.

| Model | Sum of Squares | df | F value | P-value |
|---|---|---|---|---|
| T50 ~ Elevation | | | | |
| Elevation | 263.20 | 1 | 135.31 | < 2.2E-16 |
| T50 ~ Elevation *family + (1\|family:species) | | | | |
| Elevation | 107.27 | 1 | 152.04 | < 2.2E-16 |
| Family | 11.87 | 5 | 3.37 | 0.0096 |
| Elevation:Family | 14.23 | 3 | 6.72 | 0.0002 |
| T50 ~ Elevation * species | | | | |
| Elevation | 263.20 | 1 | 467.22 | < 2.2E-16 |
| Species | 657.60 | 60 | 19.46 | < 2.2E-16 |
| Elevation:species | 88.39 | 28 | 5.60 | 2.41E-16 |

**Table S3.** Estimated marginal means of linear trends.

| Model | Intercept | Slope | Standard Error | df | Lower.CL | Upper.CL |
|---|---|---|---|---|---|---|
| T50 ~ Elevation *family + (1\|family:species) | | | | | | |
| Heliconiaceae | 50.16 | 0.0020 | 0.00023 | 487.48 | 0.00159 | 0.00248 |
| Costaceae | 50.98 | 0.0005 | 0.00035 | 495.70 | - 0.00019 | 0.00120 |
| Marantaceae | 49.24 | 0.0023 | 0.00021 | 489.21 | 0.00189 | 0.00269 |
| Zingiberaceae | 50.44 | 0.0017 | 0.00026 | 481.01 | 0.00123 | 0.00223 |

**Table S4.** Estimates to show differences of $T_{50}$ along the gradient within species.

| Model | Intercept | Slope | Lower.CL | Upper.CL |
|---|---|---|---|---|
| T50 ~ Elevation * species | | | | |
| **Heliconiaceae** | | | | |
| *Heliconia atropurpurea* | 52.78 | -0.000706 | -0.002610 | 0.001199 |
| *Heliconia irrasa* | 50.33 | 0.000775 | -0.000207 | 0.001757 |
| *Heliconia latispatha* | 49.96 | 0.002749 | 0.001767 | 0.003731 |
| *Heliconia mathiasiaexvaginalis* | 48.54 | 0.004049 | 0.003239 | 0.004859 |
| *Heliconia pogonantha* | 51.88 | 0.003346 | 0.001361 | 0.005331 |
| *Heliconia psittacorum* | 52.46 | 0.001407 | 0.000425 | 0.002389 |
| *Heliconia umbrophila* | 50.07 | 0.003891 | 0.001998 | 0.005784 |
| **Zingiberaceae** | | | | |
| *Hedychium coronarium* | 49.72 | 0.002796 | 0.001662 | 0.00393 |
| *Renealmia alpinia* | 50.45 | 0.000155 | -0.001546 | 0.001856 |
| *Renealmia cernua* | 49.66 | 0.001813 | 0.001137 | 0.002489 |
| *Renealmia pluripicata* | 48.90 | 0.003492 | 0.001097 | 0.005886 |
| *Renealmia sp2* | 52.44 | -0.000391 | -0.002037 | 0.001255 |
| *Zingiber spectabile* | 51.01 | 0.002171 | 0.001037 | 0.003306 |
| **Costaceae** | | | | |
| *Costus bracteatus* | 51.10 | 0.00053 | -0.000596 | 0.001653 |
| *Costus curvibracteatus* | 50.17 | 0.00048 | -0.002071 | 0.00304 |
| *Costus laevis* | 50.84 | 0.00037 | -0.000569 | 0.001306 |

| | | | | |
|---|---|---|---|---|
| *Costus malortieanus* | 50.44 | 0.00193 | -5.879908E05 | 0.003912 |
| **Marantaceae** | | | | |
| *Calathea crotalifera* | 49.93 | 0.00225 | 0.001361 | 0.003138 |
| *Calathea lasiostachya* | 48.79 | 0.00239 | 0.001507 | 0.003264 |
| *Calathea sp1* | 48.41 | 0.00275 | 0.000848 | 0.004657 |
| *Goeppertia cleistantha* | 48.83 | 0.00216 | 0.000171 | 0.004142 |
| *Goeppertia cuneata* | 54.45 | -0.002187 | -0.004092 | -0.000282 |
| *Goeppertia gymnocarpa* | 49.14 | 0.006139 | 0.003462 | 0.008816 |
| *Goeppertia leucostachys* | 52.35 | -0.001129 | -0.002832 | 0.000575 |
| *Goeppertia micans* | 48.81 | 0.004201 | 0.003219 | 0.005183 |
| *Goeppertia plicata* | 50.46 | -0.000134 | -0.002289 | 0.00202 |
| *Hylaeanthe hoffmannii* | 48.17 | 0.000201 | -0.001503 | 0.001904 |
| *Ischnosiphon inflatus* | 49.42 | 0.002724 | 0.001742 | 0.003705 |
| *Pleiostachya leiostachya* | 49.55 | 0.003032 | 0.000623 | 0.005441 |

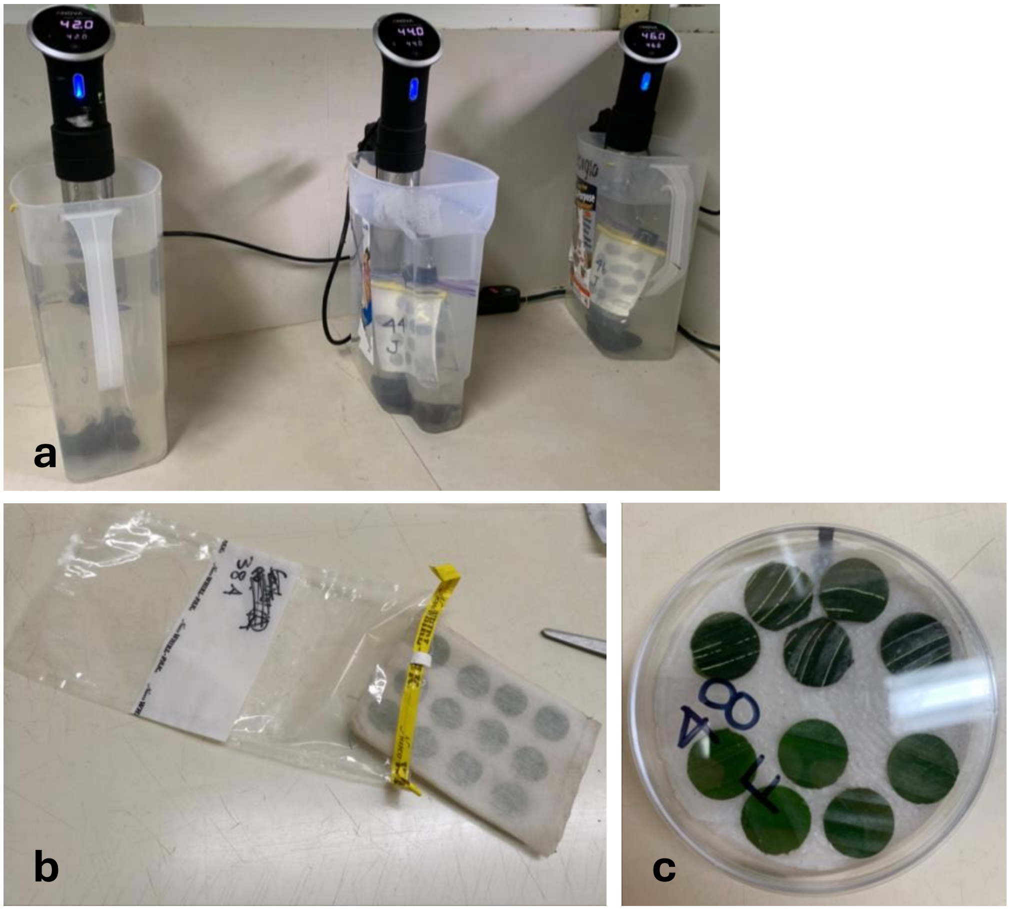


**Figure S1.** Experiment set up to determine heat tolerance ($T_{50}$). **(a)** Water baths with submerged leaf tissue placed in plastic bags. **(b)** Leaf disks in miracloth fabric and then placed in bags ready for submersion. **(c)** After temperature treatment in the water baths, leaf tissue is placed in a petri dish for 24h.

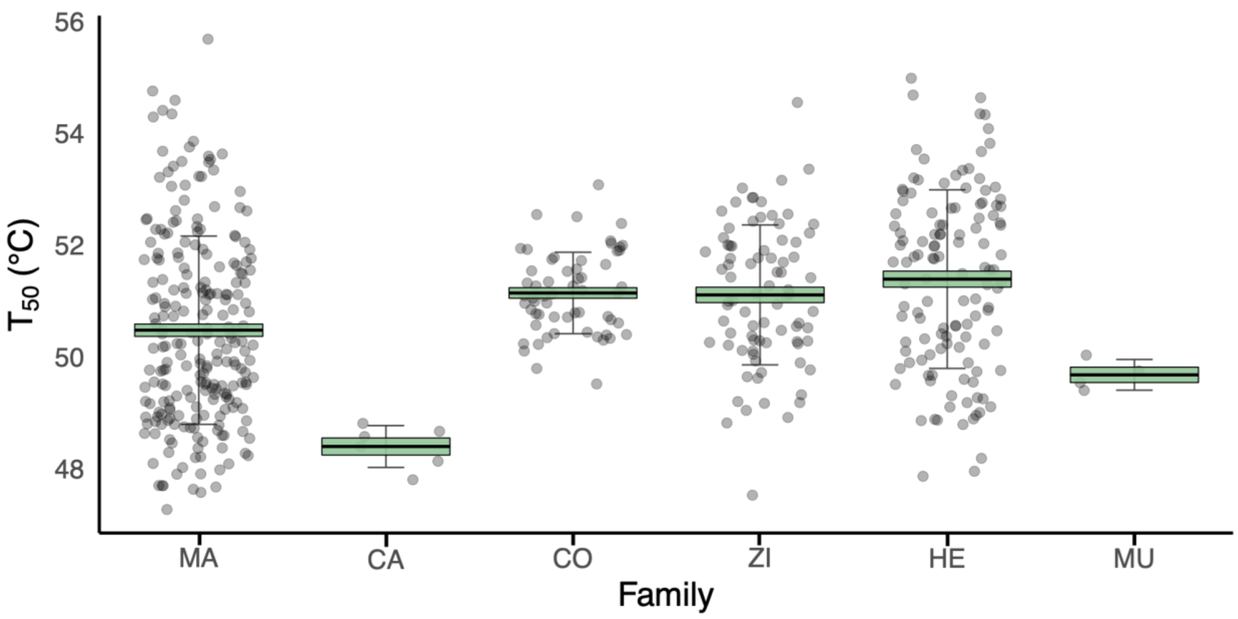


**Figure S2.** Heat tolerance of Zingiberales families along the Barva transect. Heat tolerance ($T_{50}$) was estimated as the temperature at which PSII quantum yield (fluorescence) decreases 50%. Plant families in phylogenetic order: Musaceae (MU), Heliconiaceae (HE), Cannaceae (CA), Marantaceae (MA), Costaceae (CO), and Zingiberaceae (ZI). Error bars indicate standard deviation, boxes indicate standard error, and the central line represents average heat tolerance for each family. Dots represent the mean of each species at the given family.

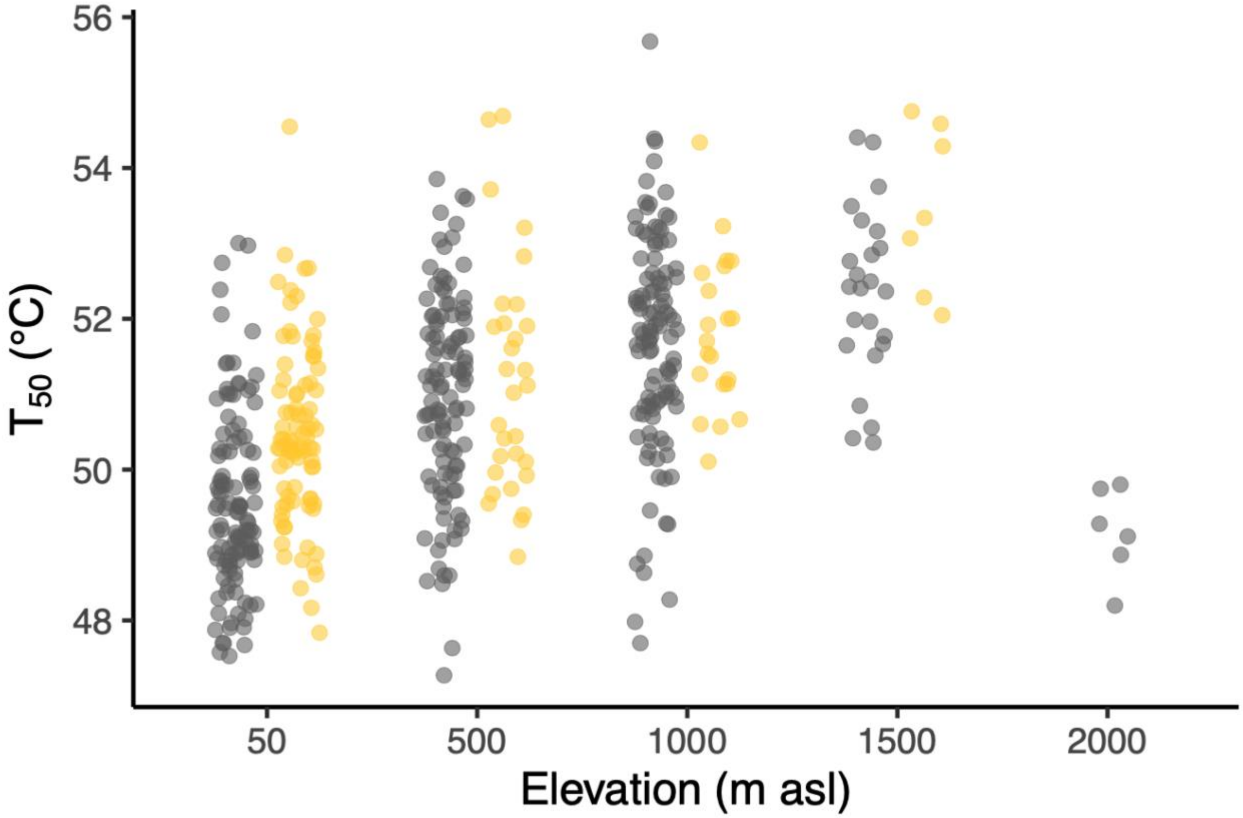


**Figure S3.** Heat tolerance ($T_{50}$) of Zingiberales individuals as a function of elevation at the Barva transect. Dots represent heat tolerance of individuals from each species at the given elevation site. Grey and yellow dots indicate if the species were collected from a shaded environment or a fully sun environment, respectively (see Table S1 for complete information on species environment).

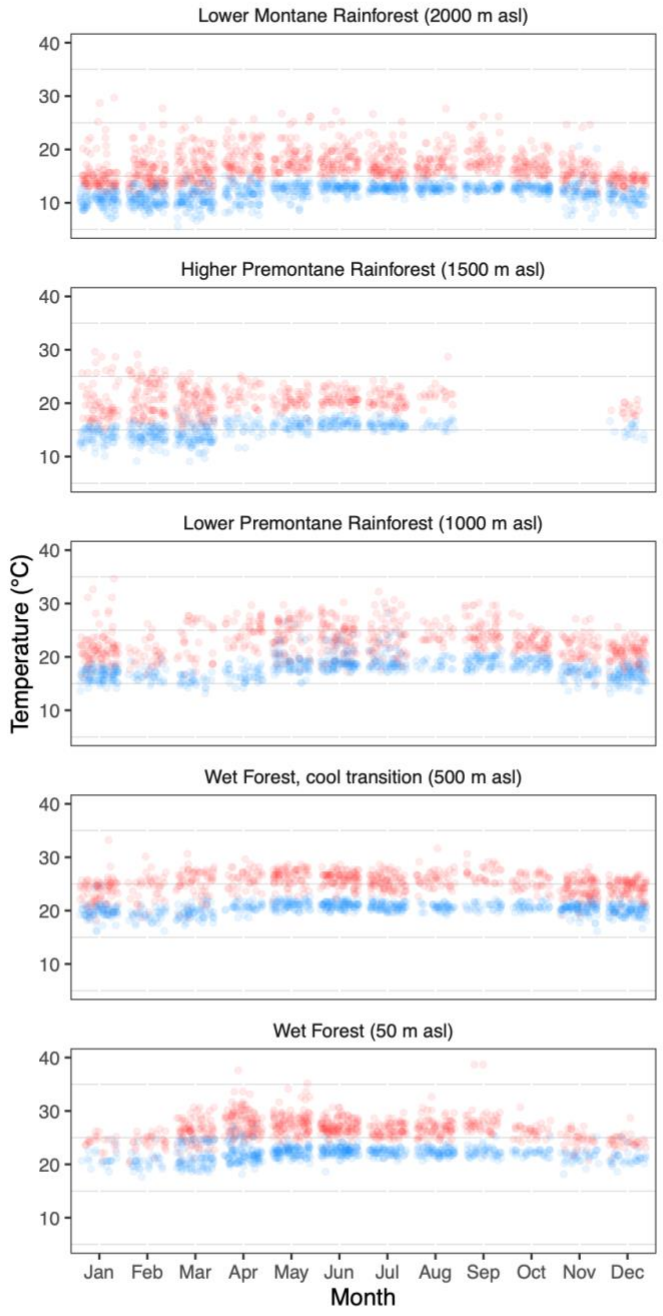


**Figure S4.** Maximum and minimum daily temperature (blue and red dots, respectively) for each elevation by month (Clark & Clark 2015) at the Barva transect, Costa Rica.

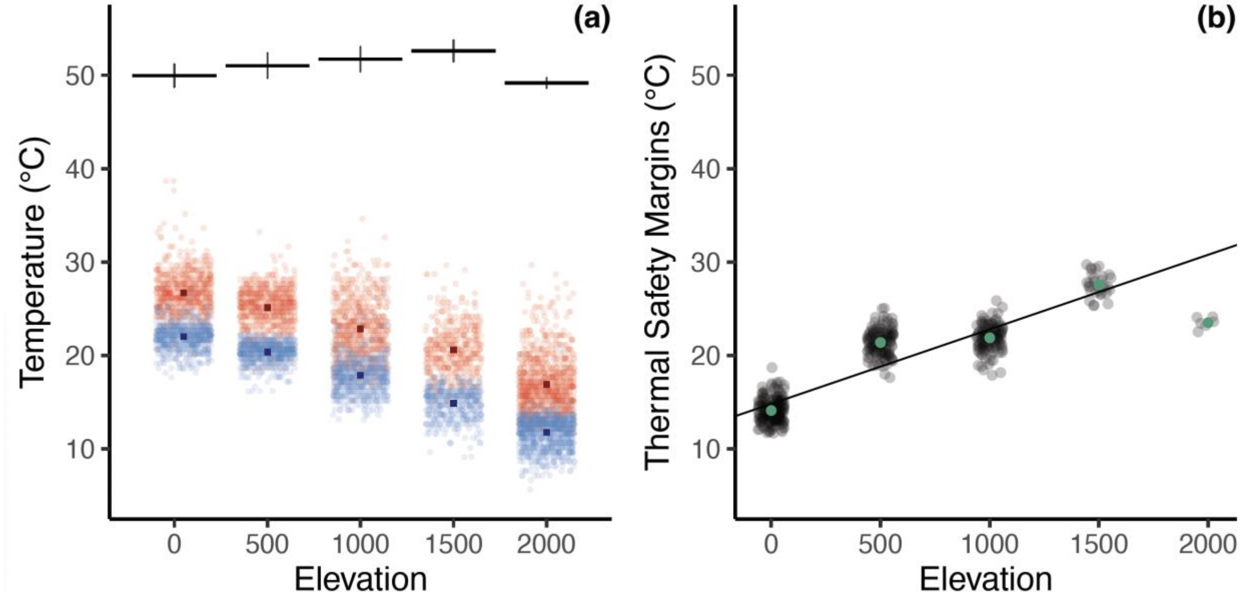


**Figure S5.** **(a)** Heat tolerances of Zingiberales along the Barva transect (black lines at the top of the figure), and maximum and minimum daily temperature (blue and red bars and dots near the bottom of the figure) for each elevation. Error bars indicate standard deviation, and the central horizontal line represents average heat tolerance for plant communities at each elevation. For the temperature data in the bottom, the central dark square represents the average for each maximum (red) and minimum (blue) daily temperature each elevation, and each dot represents the maximum and minimum daily temperature. **(b)** Thermal Safety Margins for each elevation at the Barva transect. Each dot represents TSM individual plants at each elevation. The line indicates linear regression from the model. Note that at the highest elevation only one species is present. *See* Supplementary Data Section S1for the TSM methods and statistical results.
